# Nutrient and moisture limitation reveal keystone metabolites that link switchgrass rhizosphere metabolome and microbiome dynamics

**DOI:** 10.1101/2022.06.20.496911

**Authors:** Nameer R. Baker, Kateryna Zhalnina, Mengting Yuan, Don Herman, Javier A. Ceja-Navarro, Joelle Sasse, Jacob S. Jordan, Benjamin P. Bowen, Liyou Wu, Christina Fossum, Aaron Chew, Ying Fu, Malay Saha, Jizhong Zhou, Jennifer Pett-Ridge, Trent R. Northen, Mary Firestone

**Affiliations:** Department of Environmental Science, Policy and Management, University of California, Berkeley, CA 94720; Environmental Genomics and Systems Biology Division, Lawrence Berkeley National Laboratory, Berkeley, CA 94720; Biological Systems and Engineering Division, Lawrence Berkeley National Laboratory, Berkeley, CA 94720; Institute for Biodiversity Science and Sustainability, California Academy of Sciences, San Francisco, California, USA; Plant and Microbial Biology, University of Zurich, CH-8008 Zurich, Switzerland; Department of Chemistry, University of California, Berkeley, CA 94720; University of Oklahoma, Norman, OK 73019; Noble Research Institute, Ardmore, OK 73401; Physical and Life Sciences Directorate, Lawrence Livermore National Laboratory, Livermore, CA 94550; Life and Environmental Sciences Department, University of California Merced, Merced, CA 95343

**Author notes:** co-first authors (Nameer R. Baker, Kateryna Zhalnina). **Author contributions:** M.F., J.P.-R., N.R.B., M.S., K.Z. designed research; N.R.B., K.Z., D.H., C. F., A. C., J. A. C.-N., J.S. performed research; N.R.B., K.Z., L.W., C.F., A.C., Y.F., J.S.J. generated data; N.R.B., K.Z., M.Y., J.C.-N., B.P.B., J.S. analyzed data; N.R.B., K.Z., M.Y. wrote the paper. M.F., J.P.-R., J.C.-N., J.S.J., T.R.N., J.Z., M.S. edited and finalized the manuscript.

## Abstract

Plants exude large quantities of rhizosphere metabolites that can modulate composition and activity of microbial communities in response to environmental stress. While rhizodeposition dynamics have been associated with rhizosphere microbiome succession, and may be particularly impactful in stressful conditions, specific evidence of these connections has rarely been documented. Here, we grew the bioenergy crop switchgrass (*Panicum virgatum*) in a marginal soil, under nutrient limited, moisture limited, +nitrogen (N), and +phosphorus (P) conditions, to identify links between rhizosphere chemistry, microbiome dynamics, and abiotic stressors. To characterize links between rhizosphere microbial communities and metabolites, we used 16S rRNA amplicon sequencing and LC-MS/MS-based metabolomics. We measured significant changes in rhizosphere metabolite profiles in response to abiotic stress and linked them to changes in microbial communities using network analysis. N-limitation amplified the abundance of aromatic acids, pentoses, and their derivatives in the rhizosphere, and their enhanced availability was linked to the abundance of diverse bacterial lineages from Acidobacteria, Verrucomicrobia, Planctomycetes, and Alphaproteobacteria. Conversely, N-amended conditions enhanced the availability of N-rich rhizosphere compounds, which coincided with proliferation of Actinobacteria. Treatments with contrasting N availability differed greatly in the abundance of potential keystone metabolites; serotonin, ectoine, and acetylcholine were particularly abundant in N-replete soils, while chlorogenic, cinnamic, and glucuronic acids were found in N-limited soils. Serotonin, the keystone metabolite we identified with the largest number of links to microbial taxa, significantly affected root architecture and growth of rhizosphere microorganisms, highlighting its potential to shape microbial community and mediate rhizosphere plant-microbe interactions.

**Significance:** Plants and microorganisms release metabolites that mediate rhizosphere host-microbe interactions and modulate plant adaptation to environmental stresses. However, the molecular mechanisms that underpin rhizosphere metabolite-microbiome dynamics, their functional relationships, and the biological role of plant- or microbial-produced soil metabolites remain largely unknown. Here, we found the abundances of specific classes of rhizosphere soil metabolites were responsive to abiotic stressors, and also connected to specific shifts in the rhizosphere microbial community and plant phenotypes. We propose a suite of understudied rhizosphere compounds as keystone metabolites that may structure the rhizosphere microbiome and influence plant metabolism in response to nutrient availability. These links between rhizosphere metabolites and microbial communities point to research avenues where we might leverage plant-microbe interactions to engineer enhanced rhizosphere microbiome function, plant and ecosystem health.

## Introduction

It is well-established that plants modify the chemistry and microbial communities in the rhizosphere - the soil adjacent to their roots (1-3). Rhizosphere microbial communities have increased biomass (1, 4, 5), are often less diverse than those in surrounding bulk soil (2, 3, 6, 7) and are frequently dominated by microbial taxa from specific lineages (8, 9). In addition, specific traits are enhanced in rhizosphere microbial communities, including motility (10), cell-to-cell communication or sensing (11), and nutrient uptake (12). While this indicates strong selection for specific rhizosphere competence traits, the mechanisms of rhizosphere microbial community assembly remain ill-defined. Laboratory incubations, hydroponic systems, and greenhouse studies suggest that the chemical signatures of plant exudates and mucilage - collectively termed “rhizodeposits” - are key drivers of rhizosphere microbial community structure and function. For example, as plants develop, their exudation rates and exudate chemistry change in a consistent manner (13-15) as do their rhizosphere microbial communities (1, 16). Studies conducted in hydroponic systems or with additions of exudate solutions, have observed shifts in gene expression of rhizosphere communities (17) and provide evidence that specific bacterial taxa are recruited or repelled by specific exudate compounds (18, 19). While these results support the hypothesis that root-derived chemical compounds can directly structure rhizosphere microbial communities, there have been notably few attempts to link the full diversity of the rhizosphere exometabolome to shifts in microbial community structure in complex living soils.

The rhizosphere exometabolome is a diverse chemical milieu of primary and secondary metabolites released by the plant host and rhizosphere microorganisms into the soil surrounding plant roots (12, 20). The exometabolome interacts with the soil environment (i.e., mineralogy, pH, water) and provides a varied “playground” for the cross-talk between plants and microorganisms living in soil (21-25). Plant-derived metabolites reflect plant responses to its changing environment and enable plants to modulate their metabolic interactions with microorganisms (20, 26-28), thus potentially enabling them to recruit a beneficial microbiome. At the same time, and in response, microorganisms attracted by plant-derived molecules produce rhizosphere metabolites that can alter the plant host’s phenotype and enhance its capacity to withstand environmental stresses (27, 28). Many studies characterized plant exudates in a sterile environment and demonstrate the effect of signaling metabolites on plant-microbe interactions (29-31). However, it remains unclear how the collective exometabolite chemistry of plants and microbes combines with the soil environment of the rhizosphere to impact microbiome assembly and plant adaptation to the environment.

In rhizosphere soil, diverse pools of metabolites and microbial taxa change sequentially in response to environmental drivers. This ongoing call-and-response makes it challenging to derive linear cause and effect, or the directionality of interactions. This complicates the direct assessment of functional associations between metabolites and microorganisms under different abiotic conditions. However, if changes of functionally associated metabolites and rhizosphere microbial taxa are causal, the alteration of one pool in response to changing abiotic conditions should be reflected in changes of the other. Tandem analysis of microbial communities and rhizosphere metabolite abundances should therefore be a useful approach to explore the relationships between diverse metabolites and microorganisms in soil – and a means to identify metabolite-microbiome pairs that are worthy of further experimental validation. A limited number of studies have used this approach—notably in human gut, lung, urinary tract, wastewater, and biological soil crust systems (22, 32-36) —but it has not been applied within rhizosphere soils. Linking exometabolite chemistry, microbiome assembly and plant phenotypes under abiotic stress is needed to identify functional links between specific metabolites and microbial lineages (37). These metabolite-driven changes in plant microbiomes will enable us to manipulate microbial communities and optimize plant-microbe interactions in the rhizosphere for better crop health and productivity, particularly in resource limited environments such as marginal soils.

Switchgrass (*Panicum virgatum*) is a broadly distributed tallgrass native to prairies in the Eastern and Midwestern USA (38). Drought-tolerant (39) and capable of growing in nutrient poor marginal soils (40), switchgrass is a deep-rooted perennial with significant potential to promote long-term soil carbon sequestration (41-43). These traits also make switchgrass a key candidate for cellulosic biofuel production on marginal soils that are not suitable for intensive cultivation (44). Switchgrass seedlings are susceptible to a number of biotic and abiotic stresses during the establishment phase (45), a period when mutualistic plant-microbial relationships that enhance nutrient availability, reduce moisture stress or protect against pathogens (46-49) could be critical to plant stress resilience and future yields. Switchgrass has a core microbiome of bacterial taxa that are consistently found in its rhizosphere across diverse soil and sampling environments (50-52). However, it is unclear how the switchgrass microbiome is recruited during establishment, and how abiotic stress affects these recruitment mechanisms.

In this study, we linked dynamics of switchgrass rhizosphere metabolites and microbiomes in response to abiotic stress by growing a single switchgrass genotype in a nutrient-poor marginal soil for 18 weeks under five treatments: a control, soils amended with phosphorus (+P), nitrogen (+N), both nitrogen and phosphorus (+NP), and water-limited (−W) (**Fig. 1*A***). We hypothesized that non-random co-variations in the abundances of microorganisms and switchgrass rhizosphere metabolites would emerge in response to varying abiotic stressors. Using this analysis, we identified potential ‘keystone metabolites’ - compounds that may have functional links to specific microbial lineages or abiotic stressors that significantly alter the structure of rhizosphere microbiomes. We found that N availability was the most significant determinant of both metabolite and microbial community composition, and resulted in the co-enrichment of select microbial lineages and metabolites. Keystone metabolites included compounds such as organic and aromatic acids previously linked to changes in microbiome community structure (12, 18, 19), but also N-rich compounds such as serotonin and acetylcholine that have not been investigated in the rhizosphere but are known to be strong signaling molecules in other settings (53, 54). Further, we used simplified lab experiments to show that one of the keystone metabolites identified in soil, serotonin, significantly impacts both plant phenotype and the growth of specific rhizosphere microorganisms. This study identifies keystone metabolites with unexplored potential to mediate rhizosphere communities and impact plant phenotypes under nutrient stress. In addition, our study demonstrates an approach to discovering the relationships between rhizosphere metabolites, microbial communities, plant phenotypes and abiotic stressors in complex living soils.

**Fig. 1.**
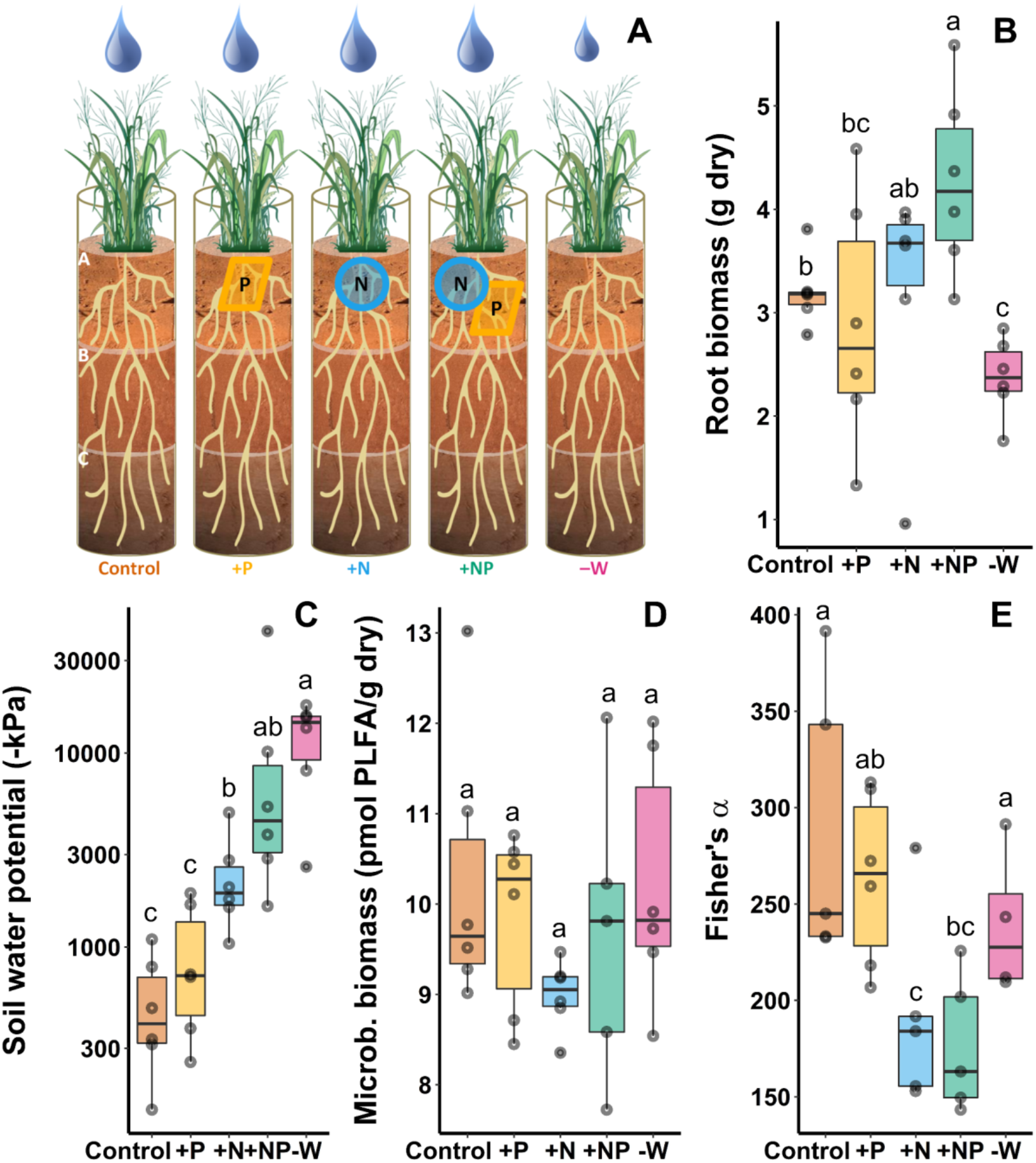
Greenhouse experiment investigating the effect of nutrient or moisture stress on switchgrass biomass, rhizosphere chemistry and microbial communities. Plants were grown in 1 meter deep mesocosms containing a marginal sandy loam soil from Anadarko OK, with recreated ‘A’, ‘B’ and ‘C’ horizons. (*A*) Schematic of experimental design illustrating five treatments: ‘Control’ with nutrient-poor marginal soil, ‘+P’, ‘+N’, and ‘+NP’ mesocosms with phosphorus and/or nitrogen amendments in the top soil horizon, and ‘-W’ mesocosms which received 50% less water relative to the other treatments. Box-whisker plots (median and 25-75% quartiles) of (*B*) switchgrass root biomass (g dry mass), (*C*) soil water potential (−kPa), (*D*) microbial biomass (pmol PLFA/g dry soil), and (*E*) microbial ɑ-diversity (Fisher’s ɑ) are shown by treatment. Letters represent significantly different post hoc pairwise comparisons via Tukey’s test (*P* < 0.05, n=6)

## Methods

### Experimental Design

Soils were collected from remnant Dust Bowl fields in Oklahoma, consisting of a nutrient-deplete Pond Creek fine sandy loam (< 0.5% total carbon, < 1 ppm total nitrogen, < 6 ppm total phosphorus) classified as a superactive, thermic Pachic Arguistoll (55). One meter deep soil mesocosms were constructed in 19.7 cm diameter impact-resistance polycarbonate tubes with the “A”, “B”, and “C” horizon soils to recreate the marginal soil environment. Six replicate mesocosms were created per treatment, for a total of 30 mesocosms. Prior to packing the mesocosms to field bulk density (1.41 ± 0.04, 1.53 ± 0.18, 1.64 ± 0.07 g dry soil cm^-3^ for the “A,” “B,” and “C” horizons, respectively), we added slow-release coated urea (ESN Smart Nitrogen, 44-0-0, Agrium) to the top horizon of the +N treatment mesocosms (0.13 g kg^-1^ soil), and rock phosphate (0-3-0, Espoma) to the +P treatment mesocosms (0.48 g kg^-1^ soil). Coated urea and rock phosphate were both added (at concentrations as above) to a third set for the +NP treatment. Two further sets of mesocosms were created - a control treatment with no nutrient amendments, and a low water (‒W) treatment which received half the water of all the other treatments once plants became established (**Fig. 1*A***). Mesocosms were watered with 50% (−W) or 100% (control, +N, +P, +NP) of the mean monthly rainfall (2012-2017, NOAA) at the field site in Oklahoma in the summer months (roughly 100 mL each day). Soil moisture sensors (EC-20; METER Group, Pullman, WA) were installed in the “A” horizon of the control and ‒W treatment to confirm differences in soil moisture. Individual clonal ramets of the Alamo switchgrass genotype, NFSG 18-01, from the Nested Association Mapping population generated at the Nobel Research Institute were planted in each mesocosm in May 2017 and grown at the University of California, Berkeley, Oxford Tract greenhouse, under a natural light regime and a 32 °C daytime and 21°C nighttime temperature cycle. After 18 weeks, each mesocosm was destructively harvested by cutting open the polycarbonate tube longitudinally, and processing the soil by horizon.

### Sample Collection and Processing

We focused our analysis of microbial communities to just the A horizon because that is the zone where the majority of root biomass and soil nutrients were found (43), and where biotic activity and nutrient exchange is presumably the greatest. Roots and associated rhizosphere soil (<2 mm from root) from the top A horizon were immediately collected for DNA extractions in 15 mL tubes containing 5 mL Lifeguard Soil Preservation Solution (QIAGEN) using chilled soil-processing trays. For metabolite extractions, roots and rhizosphere soil from all three horizons were collected and immediately placed on dry ice and then stored at -80 °C. Bulk soils (> 4 mm from roots) from the A horizon were collected and stored at 4 °C for gravimetric soil moisture measurements and all remaining roots were collected, dried and weighed. For full details on sample processing and assessments of root biomass and bulk soil pH water potential, and phospholipid fatty acid analysis (PLFA) microbial biomass, see Sher, Baker et al. 2020 (43).

### DNA Extraction and Sequencing

Additional details for the rhizosphere soil DNA extraction, metabolite extraction and analysis, sequencing, bioinformatics, and statistical methods are provided in the Supplemental Materials and Methods. Briefly, rhizosphere soil in Lifeguard solution was centrifuged to pellet and DNA was extracted from single 0.5 g aliquots using a modified RNA/DNA phenol chloroform co-extraction protocol via bead-beating (56, 57). Notably, the 5% hexadecyl-trimethyammonium bromide/0.7 M NaCl/240 mM K-PO_4_ buffer (pH 8) was modified to include 1% β-mercaptoethanol, PEG 8000 was used in place of PEG 6000, and GlycoBlue was used to stain nucleic acid pellets. Microbial community composition was characterized with a sequencing library prepared at the University of Oklahoma via a phasing amplification technique targeting the V4 region of the 16s ribosomal RNA gene with the 515F and 806R primer set (58, 59). Samples were sequenced on the Illumina MiSeq platform with 2×250 bp format.

### Bioinformatics

All initial bioinformatics processing and production of amplicon sequence variants (ASVs) by DADA2 (60) were conducted within Qiime2 (61), with taxonomy assigned via the SILVA database (release 132) (62). 10,516,421 raw reads were processed into 7856 denoised ASVs that accounted for 3,942,210 reads. Subsequent processing, visualization, and statistical tests of sequence data were performed in R version 3.6.0 (R Core Team, 2020), primarily within the phyloseq package (63). Chloroplast, mitochondrial, and bacterial or archaeal sequences that lacked designation at the phylum level were discarded, leaving 7481 ASVs accounting for 3,792,761 reads. Singletons and doubletons were removed for all analyses other than ɑ-diversity, resulting in 7093 ASVs accounting for 3,792,087 reads (104,776-194,526 per sample). Differentially abundant ASVs between each of the individual treatments and the control samples in the top horizon were determined with the DESeq2 package (64), and differentially abundant ASVs whose responses to a treatment were driven by only one sample were removed from subsequent analyses. Analyses of β-diversity were performed via permutational analysis of variance (PERMANOVA) on rarefied sets of 100,000 reads per sample using Unifrac distance matrices.

### Metabolite Extraction and Analysis

Soil metabolites were extracted from the “A”, “B”, and “C” soil horizons using a method described in Swenson et al. (2015) (65). Briefly, rhizosphere soil samples were shaken in ice-cold liquid chromatography-mass spectrometry (LC-MS)-grade water at 200 rpm for 1 h at 4 °C and centrifuged at 3220 g for 15 min at 4 °C. The supernatant was filtered through 0.45 μm syringe filters (Pall Acrodisc Supor membrane) and lyophilized. Lyophilized extracts were resuspended in 100% methanol with internal standards, filtered using 0.22 μm microcentrifuge PVDF filters (Merck Millipore), and aliquots of 150 μl of methanol extracts were analyzed using normal-phase LC-MS with a HILIC-Z column and an Agilent 1290 LC stack. MS and MS/MS data were collected using a Q Exactive Orbitrap MS (Thermo Scientific) (see Supplemental Materials and Methods for additional details).

Metabolomics data were analyzed using Metabolite Atlas software to obtain extracted ion chromatograms and peak heights for each metabolite (66). Metabolite identifications were verified with authentic chemical standards based on matching m/z better than 5 ppm for positive mode, 15 ppm for negative mode, retention time difference ≤ 0.5 min, and/or MS/MS fragment matching score of > 0.6 as calculated by the Stein and Scott ‘composite’ algorithm with modifications (67) (**Table S4**). As defined by the Metabolomics Standards Initiative (68), any two of these orthogonal measures supports a level 1 identification for the identified metabolites (provided that the third measure did not invalidate the identification). A total of 100 level 1 unique metabolites were identified, and seven metabolites were classified as ‘unresolvable” metabolites’ due to structural isomers (**Table S4**). All identified metabolites were detected in at least four out of six replicates from at least one treatment.

Significant differences in switchgrass rhizosphere metabolite profiles in response to the five nutrient and water stress treatments and between the three soil horizons were determined with PERMANOVA. The magnitude of change (Δ of metabolite abundance) for all significantly changed metabolites (*P* < 0.05) was calculated by scaling metabolite peak heights from 0-1, where “1” is the highest peak height of each metabolite across all samples, and then subtracting the scaled metabolite abundances observed in nutrient-depleted marginal soil (C) from the treatments where N (+N; +NP) or P (+P) had been added or where water was limited (−W). A positive Δ indicates an increase in metabolite abundance in a specific treatment (**Fig. 2*A,C,E***).

**Fig. 2.**
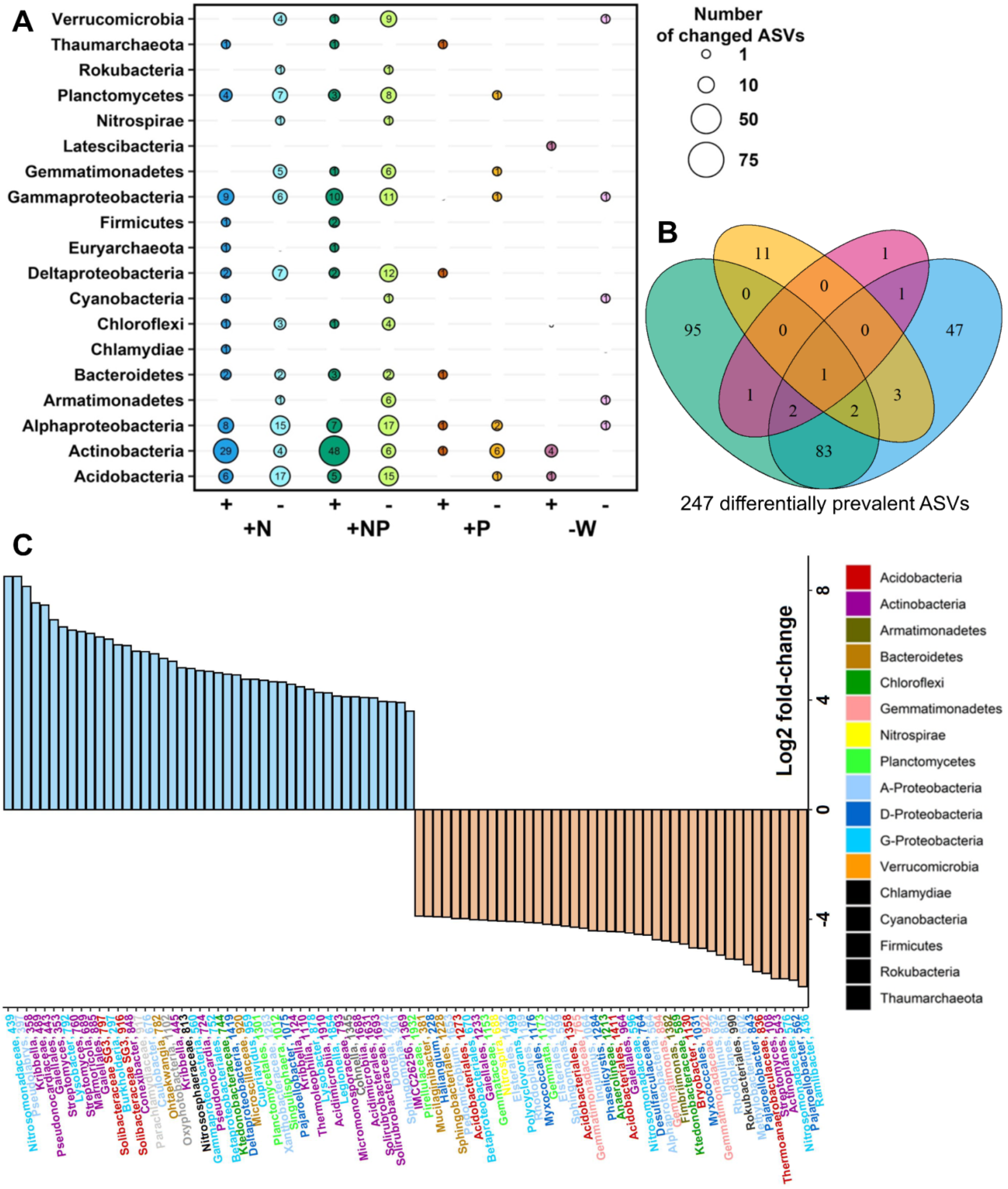
Influence of nutrient and water limitation on switchgrass rhizosphere microbial community structure assessed by DESeq2 analysis (adjusted *P* < 0.01, see **Supplementary Information** for details). (*A*) Number of positively (+) and negatively (−) responsive amplicon sequence variants (ASVs) in nutrient amended and water-limited treatments (+N, +NP, +P, -W) as compared to control soils, arranged by phyla (bubble size reflects the number of responsive ASVs). Empty cells indicate no responsive ASVs from that phylum. (*B*) Number of unique and shared amplicon sequence variants (ASVs) that changed in prevalence in response to each treatment relative to controls. (*C*) The top-50 ASVs that increased (+ Log2 fold-change) versus decreased in prevalence (− Log2 fold-change) in response to the +N treatment. ASVs are presented at the highest available taxonomic resolution, and are colored by class for *Proteobacteria* and by phylum for all other phyla.

### Rhizosphere Metabolite - Microbiome Associations

To determine covariance between metabolites and microbial ASVs, we constructed a correlation network based on the relative abundances of metabolites and relative abundances of ASVs across all treatments. To prevent false positives, we only included ASVs with non-zero abundances in at least 15 of the 25 samples. Spearman correlations were calculated for each of the metabolite-metabolite, ASV-ASV and metabolite-ASV pairs. A Random Matrix Theory (RMT)-based approach determined 0.710 as the correlation coefficient cutoff that controls false discovery in our network by separating noise vs. non-random correlations (69). This RMT-based approach has been previously used to construct correlation-based networks of complex microbial systems (6, 70), and is available through the Molecular Ecological Network Analysis (MENA) pipeline (http://ieg4.rccc.ou.edu/MENA/). To construct the networks, we included only metabolite-ASV pairs with an abundance correlation coefficient above the threshold of 0.710, and discarded links within metabolite species or within 16S ASVs to construct the network. Discarding non-metabolite-ASV links partly alleviates the potential bias caused by compositional data, as the abundances of ASVs and metabolites were independently derived; also, compositional data bias should be a minor problem in high-diversity communities (71). Positive and negative correlations correspond to positive and negative links, respectively.

Based on previously developed criteria (70, 72, 73), we defined putative ‘keystone metabolites’ or ‘keystone ASVs’ from an individual node’s role in network topology as follows: the network was separated into modules using a fast greedy algorithm, and the within-module connectivity (zi) and among-module connectivity (pi) were calculated for each node (72). Nodes with zi >2.5 are designated as module hubs, while nodes with pi >0.62 were designated as connectors among different modules. Nodes with both zi >2.5 and pi > 0.62 stretch among the whole network were designated as network hubs (73). These module hubs and network hubs were defined as keystone metabolites or keystone ASVs based on the node’s identity. Correlation calculations, network construction, and network topology analysis were conducted with the igraph package (74). The network was visualized using Cytoscape (75). In addition, to identify associations between metabolites, microbial communities, and treatments, we performed hierarchical clustering analysis using the vegan package (76). For the analysis we selected differentially abundant ASVs determined with the DESeq2, that had more than three significant positive or negative Spearman’s rank correlations with metabolites (*r* > 0.7, *P* < 0.05) and metabolites with more than one significant positive or negative correlation with ASVs (*r* > 0.7, *P* < 0.05).

### Plant phenotype response to serotonin

To test the effect of serotonin on plant growth, surface-sterilized Alamo switchgrass seeds were sown on ⅕ Murashige and Skoog (MS) basal salt mixture M524 (Phyto Technology Laboratories) (0.87 g/L MS salts, pH 7.3, and 8 g/L agar). Nine biological replicates (n = 9) of seven-day-old seedlings were transferred to ⅕ MS agar plates supplemented with 0.1 mM serotonin (Sigma-Aldrich) or with purified H_2_O. Plates with seedlings were incubated at 24 °C on a 16 h/8 h day/night cycle, with humidity maintained at 70% and irradiance at 250 μE m^−2^ s^−1^. After twenty-five days, switchgrass plants were harvested, and root and shoot biomass were measured. Root length and root number were quantified using the SmartRoot plugin (version 4.21) in ImageJ (version 2.0.0) (77). Significant differences between treatments were determined using an ANOVA test (*P* < 0.05).

### Microbial response to serotonin

To test the effect of serotonin on microbial growth responses, we selected eight bacterial isolates from a suite previously cultured from Oklahoma marginal soils planted with switchgrass. We mapped 16S rRNA gene sequences from these isolates to the 16S amplicon sequences from the switchgrass rhizosphere in this study to ensure selected isolates were closely related to those observed in the rhizosphere. Isolates and observed ASVs were determined to have closely related V4 regions of their 16S rRNA genes (extracted using BLASTN) if their E values were <1 × 10^−10^ and matched ≥97% of gene sequence homology. These isolates were then assigned the serotonin response pattern (positively- or negatively-correlated) corresponding to the ASV they were closely related to in this study. Eight isolates (four with putative positive correlations with serotonin and four with negative) were selected to analyze their growth response to serotonin-spiked growth medium. These isolates represent common rhizosphere genera: *Bradyrhizobium, Reyranella, Mucilaginibacter, Methylobacterium, Shinella, Paenarthrobacter, Burkholderia*, and *Mesorhizobium*. Growth curves were performed in 1/10 R2A medium with either 0, 0.1, or 0.5 mM serotonin. Four replicates of each isolate were inoculated in a 96-well plate and grown at 30 °C, shaking once per hour at 200 rpm before optical density measurement at 600 nm (OD_600_). After 130 hours of isolate growth with 0.1 or 0.5 mM serotonin, the culture OD_600_ was compared to that of a control treatment without serotonin (0 mM). Optical density responses were analyzed using a Kruskal-Wallis test after the OD_600_ of uninoculated blanks was subtracted from the inoculated samples.

## Results

### Plant and soil responses to nutrient and water treatments in the “A” horizon

Switchgrass root biomass in the “A” horizon varied significantly by treatment (*P* < 0.001, **Fig. 1*B***) and was highest in the +NP treatment (4.80 ± 1.04 g; mean ± SD) and lowest in the -W treatment (2.45 ± 0.24 g). The soil water potential at harvest also varied significantly by treatment (*P* < 0.001, **Fig. 1*C***); the control soils were the wettest (−527 ± 352 kPa), treatments with higher root biomass (+N, +NP) had generally drier soils, and the -W treatment had the driest soils (−12,100 ± 5700 kPa). Soil microbial biomass, measured by PLFA, did not vary significantly by treatment (**Fig. 1*D***), with an average 9.81 ± 1.25 pmol PLFA / g dry soil observed across all treatments.

### Microbial diversity in rhizosphere soil

To evaluate how nutrient availability and moisture stress affected microbial diversity in the rhizosphere of switchgrass, we analyzed 7841 amplicon sequence variants (ASVs) from the uppermost soil horizon. Microbial ɑ-diversity varied significantly by treatment according to a suite of metrics including Fisher’s ɑ (*P* = 0.01, **Fig. 1*E***), the Chao1 index (*P* = 0.009), the Shannon index (*P* < 0.001), and the Simpson index (*P* = 0.002) (**Fig. S1**). Phylogenetic diversity also varied significantly by treatment (*P* = 0.01), but mean pairwise distance between ASVs did not. In general, ɑ-diversity metrics were significantly higher in controls and in the +P treatment relative to soils where N was added, while reduced watering did not have a significant effect relative to controls (**Fig. S1**).

At the phylum level, rhizosphere communities were dominated by *Actinobacteria* and *Proteobacteria*; together, these phyla made up 46.4 ± 8.2 % and 26.5 ± 2.3 % of the sequences observed in each sample, respectively (**Table S1**). *Acidobacteria* and *Verrucomicrobia* were the only other phyla that comprised >4 % of the community in each sample, on average. Other phyla that comprised >1 % of the average community were (in order of abundance) *Firmicutes, Chloroflexi, Gemmatimonadetes, Bacteroidetes*, and *Planctomycetes* (**Fig S2*A***). The phyla *Actinobacteria* and *Acidobacteria* were the only dominant phyla (>4% relative abundance) that varied significantly between treatments (*P* < 0.05).

Community β-diversity varied significantly by treatment (*P* < 0.001) according to Unifrac distances, with the strongest differences resulting from N-addition (**Fig. S2*B***). Communities from the +N and +NP treatments were significantly different (*P* < 0.05) from those in the control and +P treatment, and communities from the -W treatment were significantly different from those in the control and +NP treatment (**Table S2**).

### Impact of nutrient and moisture limitation on rhizosphere ASVs

To assess the effects of abiotic stressors on specific ASVs and determine which ASVs were driving community-level differences between treatments, we used DESeq2 to identify lineages that were differentially abundant in the +N, +P, +NP, and -W treatments relative to the control (**Fig. 2*A***). N addition (+N, +NP) caused the strongest shifts in community composition (**Fig. 2*A***), with the abundance of 247 ASVs significantly affected by treatment: 184 by the +NP treatment, 139 by +N, 17 by +P, and 6 by -W. While there was some overlap in the effect of treatments on ASVs (**Fig. 2*B***), many ASVs were solely affected by the +NP and +N treatments (95 and 47, respectively). Notably, 83 ASVs were uniquely affected in both the +N and +NP treatment soils, evidence of a strong N addition effect. Relatively few ASVs were solely affected by the +P and - W treatments (11 and 1, respectively). DESeq responsive ASVs and their taxonomy are listed in **Table S3**.

We defined ASVs that were more likely to be found (or not found) in a given treatment as ‘positive’ (or ‘negative’) responders. The taxonomic identity of ASVs that responded to the +N or +NP treatments depended on whether they were positive- (increased in abundance) or negative-responders (decreased in abundance). Positive-responders to N-application (+N or +NP) were dominated (>50%) by ASVs from the phyla Actinobacteria (e.g., *Kribella, Streptomyces, Marmoricola, Conexibacter*) (**Fig. 2*C***). ASVs with negative-responses to N were more taxonomically diverse as a group. Negative-responders came from >17 classes, with the majority belonging to the Acidobacteria (e.g., *Bryobacter, Acidobacteriales, Acidobacteria SG17)*, classes Alphaproteobacteria (e.g., *Elsterales, Xanthobacteraceae, Methylobacterium*), Deltaproteobacteria (e.g., *Pajaroellobacter*, other *Myxococcales*), Gammaproteobacteria (e.g., *Burkholderiaceae, Nitrosomonadaceae*), the Planctomycetes (e.g., *Gemmataceae*), and the Verrucomicrobia (e.g., *Pedosphaeraceae*)(**Fig. 2*C***).

Far fewer ASVs responded to the +P and -W treatments (**Fig. S3*A,B*** respectively) and there were few taxonomic patterns. Of 17 responders to the +P treatment, only 5 ASVs increased in abundance in response to P amendment, all from different phyla. A majority of the ASVs (n=5) that decreased in response to P addition belonged to *Actinobacteria*. The -W treatment had the fewest ASVs change in abundance relative to the control soil, but 6 ASVs responded positively to water limitation, with the response driven by five ASVs from *Actinobacteria*.

### Treatment-induced changes in rhizosphere metabolite chemistry

LC-MS-based metabolomics was used to identify rhizosphere metabolites under different nutrient additions and water limitation. 100 metabolites were identified across five treatments and three soil horizons (**Table S4**). Analysis of rhizosphere metabolites revealed compositional changes in all treatments, with the +N and +NP treatments exhibiting the greatest differences compared to the nutrient-poor control soil (**Fig. 3**). Across all soil horizons seventeen metabolites were significantly more abundant (PERMANOVA: *P* < 0.05) in the rhizosphere of the marginal control soils when compared to nitrogen (+N and/or +NP) supplemented treatments (**Fig. 3*A,B*, Fig. S4**), and declined in abundance when nitrogen was added. The majority of these 17 metabolites were organic acids (n=12) and about half contained an aromatic ring (*n*=9); the remaining metabolites included pentoses and pentose alcohols (n=3), vitamin B and a lactone. Conversely, addition of N (+N and/or +NP) significantly increased the abundance of 35 N-containing rhizosphere metabolites and one sugar for all three soil horizons. This included amino acids, nucleosides, and quaternary amines, as well as N-containing azoles such as allantoin and N-containing indoles such as serotonin (**Fig. 3*C,D*, Fig. S4**). Moisture stress also had a significant effect on an array of rhizosphere metabolites; 17 metabolites increased in abundance in response to the -W treatment, including a number that are known osmolytes (amino acids, quaternary amines, sugars) (**Fig. 3*E,F*, Fig. S4**). P amendment had a much smaller effect on rhizosphere metabolite chemistry than the other treatments; in the +P treatment, only seven metabolites significantly changed in abundance relative to the control (**Fig. S4**).

**Fig. 3.**
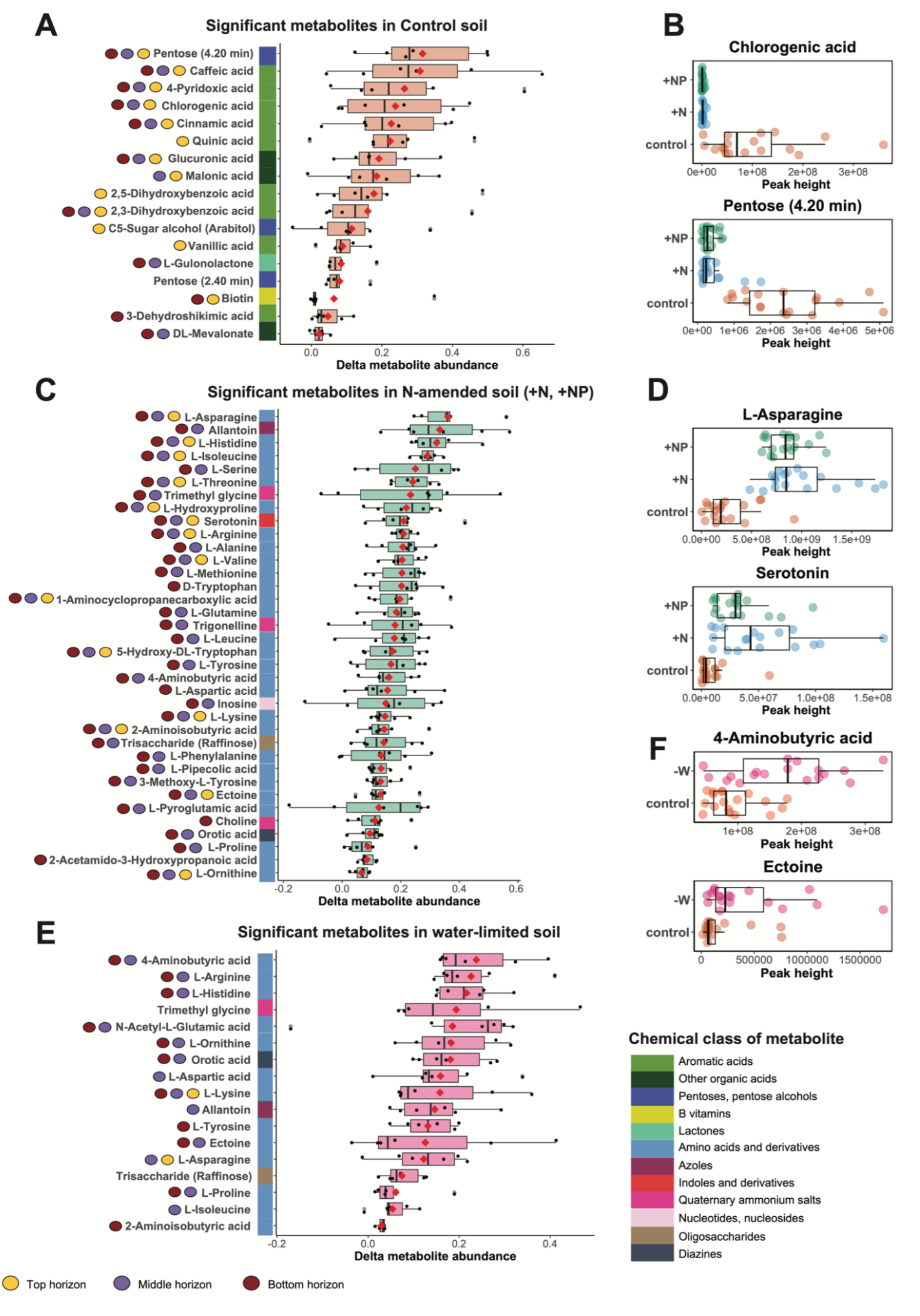
Significant changes in switchgrass rhizosphere metabolite profiles in response to five nutrient and water stress treatments (n replicates=6), assessed by PERMANOVA (*P* < 0.05, see **Supplementary Information** for details). (*A*) Metabolites significantly enriched (*P* < 0.05) in a nutrient-depleted marginal soil (control) compared to treatments where N was added (+N; +NP). (*C*) Metabolites that increased (*P* < 0.05) in abundance in response to N addition (+N, +NP) compared to the control soil. (*E*) Metabolites that increased in abundance (*P* < 0.05) in response to water limitation (−W) compared to the control soil. Y-axis circles next to each metabolite represent the soil horizons where the metabolite had a significantly different abundance. Unresolvable metabolites are indicated by parentheses (22, 36). The red diamond inside each box denotes the mean and the horizontal line denotes the median. (*B, D, F*) Box-whisker plots of the abundance of example metabolites are shown to illustrate treatment effects across all three horizons. Points reflect a single metabolite per sample, the outer boxes indicate the first, second and third data quartiles, and whiskers indicate the range of the points excluding outliers.

We also analyzed metabolite changes in response to the treatments for each soil horizon. The top-soil horizon responded the most to nutrient limitation, with 13 metabolites increased in abundance when N was limited out of the 17 metabolites that changed across all horizons (**Fig. 3*A***). Nitrogen addition resulted in the most significant changes in metabolite abundances of any treatment in the middle and bottom horizons (**Fig. 3*B***), where nearly any metabolite with a significant response to N addition was found to increase in abundance, and very few were observed to decrease. Notably, the bottom and middle soil horizons revealed more profound metabolite responses to water limitation than the top horizon, where only two out of 17 metabolites increased (**Fig. 3*E***).

### Associations between metabolites, microbial ASVs and abiotic stresses

To identify relationships between microbes and metabolites, we used Spearman’s rank correlations and hierarchical clustering of differentially abundant rhizosphere ASVs defined by the DESeq analysis and metabolites observed in the “A” soil horizon. This analysis groups treatment-responsive rhizosphere metabolites and ASVs by their degree of correlation, to identify clusters with similar behavior. Hierarchical clustering of the most responsive ASVs and differentially abundant metabolites revealed two large microbial-metabolite clusters. Cluster #1 (**Fig. 4**) contains microbial ASVs (n=8) and rhizosphere metabolites (n=17) that increased in abundance in the N-amended treatments (+N, +NP), including ASVs from the *Actinobacteria* (*Paenarthrobacter, Solirubrobacter*), *Alphaproteobacteria* (*Sphingomonas, Pseudolabrys*), *Gammaproteobacteria* (*Nitrosomonadaceae*) and metabolites with N-rich compounds (amino acids, azoles, quaternary amines). Cluster #2 (**Fig. 4**) includes metabolites (n=8) and microbial ASVs (n=29) with higher relative abundance in the unamended control soils. ASVs in this cluster were distinct from the ASVs identified in the Cluster #1 and include diverse microbial classes from the *Alphaproteobacteria* (*Rhodoplanes, Acetobacteraceae*), *Deltaproteobacteria* (*Desulfarculaceae, Myxococcales*), *Verrucomicrobia* (*Pedosphaeraceae, Chthoniobacter*), *Acidobacteria* (*Bryobacter, Unclassified Acidobacteria*), as well as ASVs from *Planctomycetes, Nitrospirae, Armatimonadetes, Gemmatimonadetes, Bacteroidetes* and *Actinobacteria*. The majority (75%) of rhizosphere metabolites that co-varied with the ASVs from the Cluster #2 were organic acids, particularly aromatic acids (chlorogenic, cinnamic, caffeic, 4-pyridoxic, 2,3-dihydroxybenzoic acid).

**Fig. 4.**
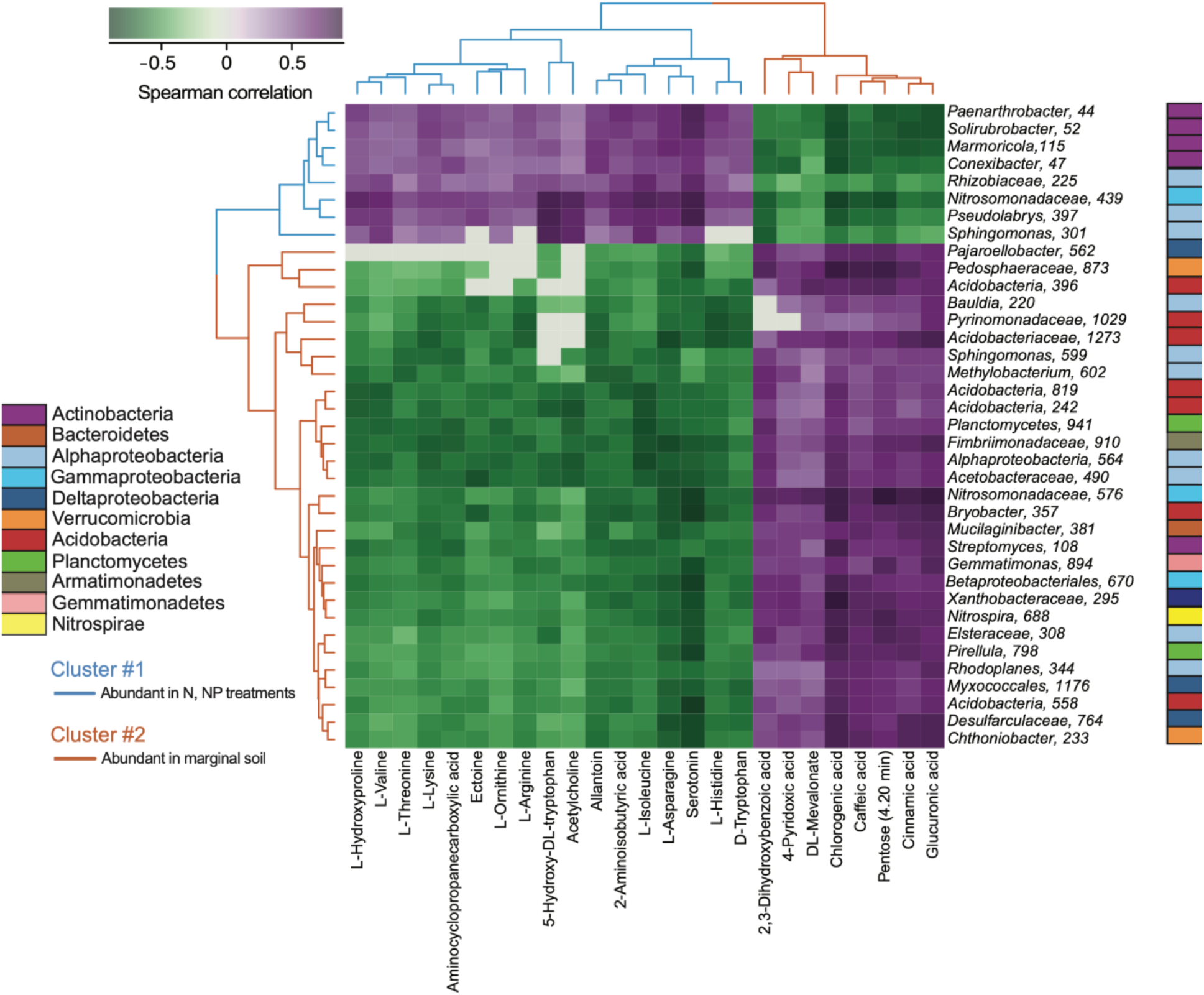
Heatmap representing the top co-varying microbial taxa and metabolites in the rhizosphere of switchgrass grown with five soil nutrient and water treatments. Top associations between metabolites (columns) and ASVs (row) include (i) DESeq2-determined differentially abundant ASVs (n=37) with more than three significant positive or negative correlations (Spearman’s rank correlation, *r* > 0.7, *P* < 0.05) with metabolites; and (ii) metabolites (n=25) with more than one significant positive or negative correlation (Spearman’s rank correlation, *r* > 0.7, *P* < 0.05) with ASVs. Hierarchical clustering shows two clusters of metabolite-ASV correlations. Cluster #1 (blue lines) represents metabolites and ASVs that were more abundant in the rhizosphere when nitrogen was added (+N, +NP treatments) and Cluster #2 (brown lines) includes metabolites and ASVs that were more abundant in nitrogen-poor marginal soil (controls). Purple colors in the heatmap represent positive Spearman correlations, white represents no correlation, and green colors represent negative correlations between metabolites and ASVs.

### Metabolite-microbial rhizosphere community network

In our bipartite co-occurrence network of rhizosphere ASVs and metabolites (**Fig. 5*A***), 117 ASVs connect to 31 metabolites via 368 links, including 153 positive and 215 negative links, with an average of 5 links per node (**Table S5**). We identified five module hubs and one network hub as putative “keystone metabolites” (**Table S5, Fig. 5*B***). Three metabolites reflect modules dominated by **negative** correlations, including serotonin, acetylcholine, and ectoine (modules 2, 3, and 4, respectively), with serotonin exhibiting negative links (83%) with a wide range of bacteria, and positive links primarily with *Actinobacteria* ASVs. Module 1, the largest, was driven by **positive** interactions with chlorogenic acid, glucuronic acid, and cinnamic acid, and included 78% positive links with bacterial ASVs from a diverse range of lineages and negative links primarily with ASVs from *Actinobacteria*. There was no module hub observed for Module 5, which was dominated by metabolite nodes instead of ASVs.

**Fig. 5.**
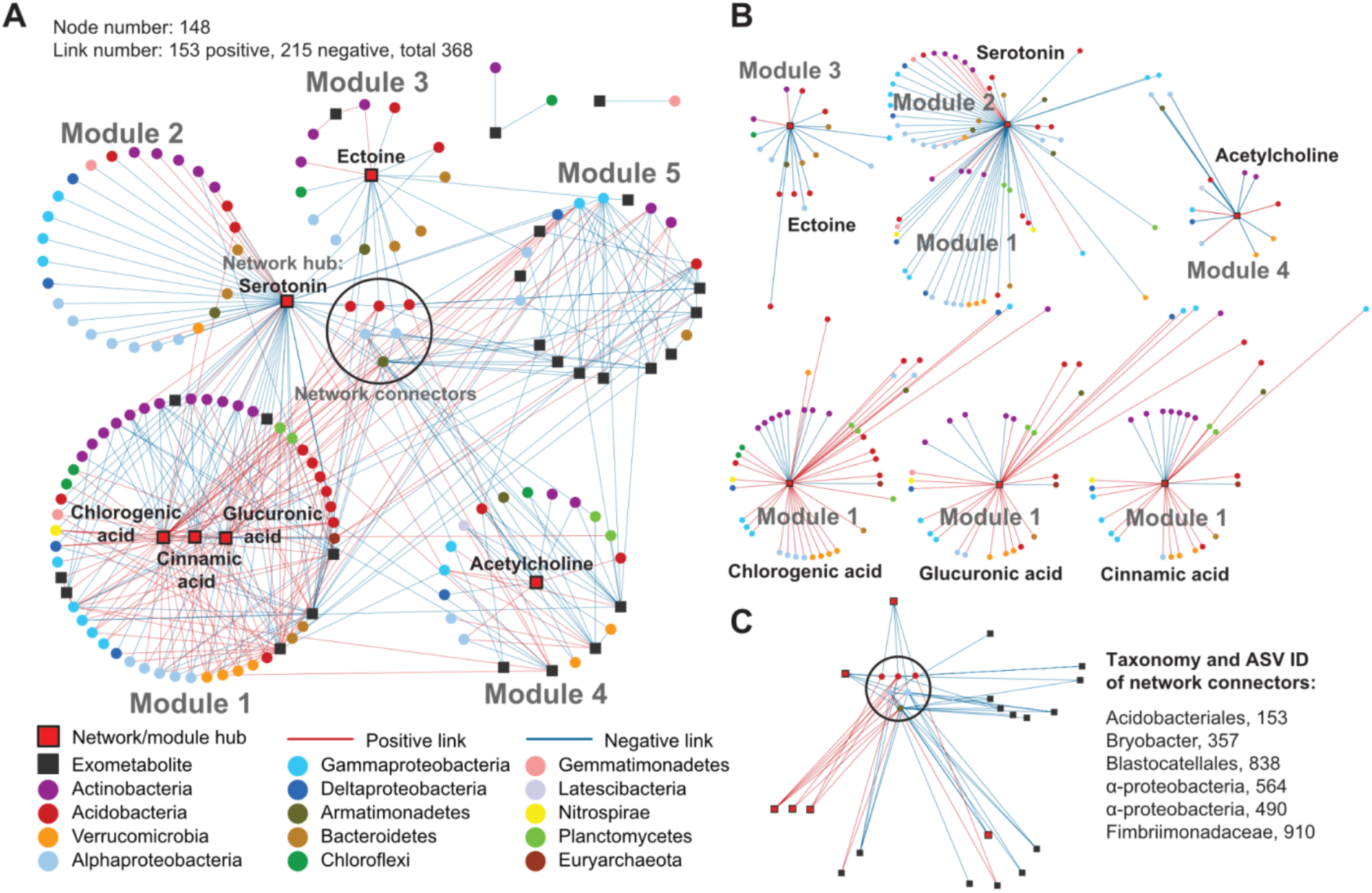
Co-occurrence network of switchgrass rhizosphere metabolites and microbial ASVs exposed to five soil treatments in a greenhouse study. (*A*) An association network between 908 16S ASVs and 100 rhizosphere metabolites. Nodes with circle symbols represent 16S ASVs, and nodes with square symbols represent metabolites. Links between nodes are based on Spearman correlations (*r* > 0.710) of their relative abundances, red for positive correlation and blue for negative correlation. There are a total of 148 nodes and 368 links in this network. The network separates into five major modules, or highly connected groups of nodes, shown as the five numbered circles. Red filled squares highlight rhizosphere metabolites that act as network and module hubs, which are the nodes with dense connections to other nodes within the entire network (network hub) or a module (module hub). The six microbial ASV nodes at the center serve as connectors of different modules, or the nodes linking different modules. (*B*) Subnetworks of rhizosphere metabolites that formed module hubs and their neighboring microbial nodes. (*C*) Subnetworks of microbial nodes that serve as connectors, and their linked rhizosphere metabolite. Microbial ASVs are colored by class for *Proteobacteria* and by phylum for all other phyla.

The six connector ASVs behaved similarly to one another (**Fig. 5*C***), forming most of their positive links (4/5) with organic acids - including the three aforementioned module hubs and 2,3-dihydroxybenzoic acid in Module 4 - and forming most of their negative links (10/12) with non-organic acid metabolites. Approximately half of the ASVs in the network were also identified as differentially abundant by DESeq (**Table S3**), and nearly all of these were responsive to the +N or +NP treatment but not the -W or +P treatments.

### Serotonin impacts on plant biomass and rhizosphere isolates

Serotonin was identified as a key hub metabolite in the reconstructed network (**Fig. 5**); it formed the largest number of ASV links and had the largest number of significant ASV correlations (**Fig. 4** and **Fig. 5**). Given serotonin’s known role in gut bacterial-host interactions (54) and phenotypic effects on *Arabidopsis* (78) we conducted a follow-up study to examine its effect in the switchgrass rhizosphere. Switchgrass seedlings grown with 0.1 mM serotonin had increased root and shoot biomass (**Fig. S5**), promoted the number of secondary roots (**Fig. 6*A***), and increased secondary root length (**Fig. 6*B***) (*P* < 0.05, n=9).

**Fig. 6.**
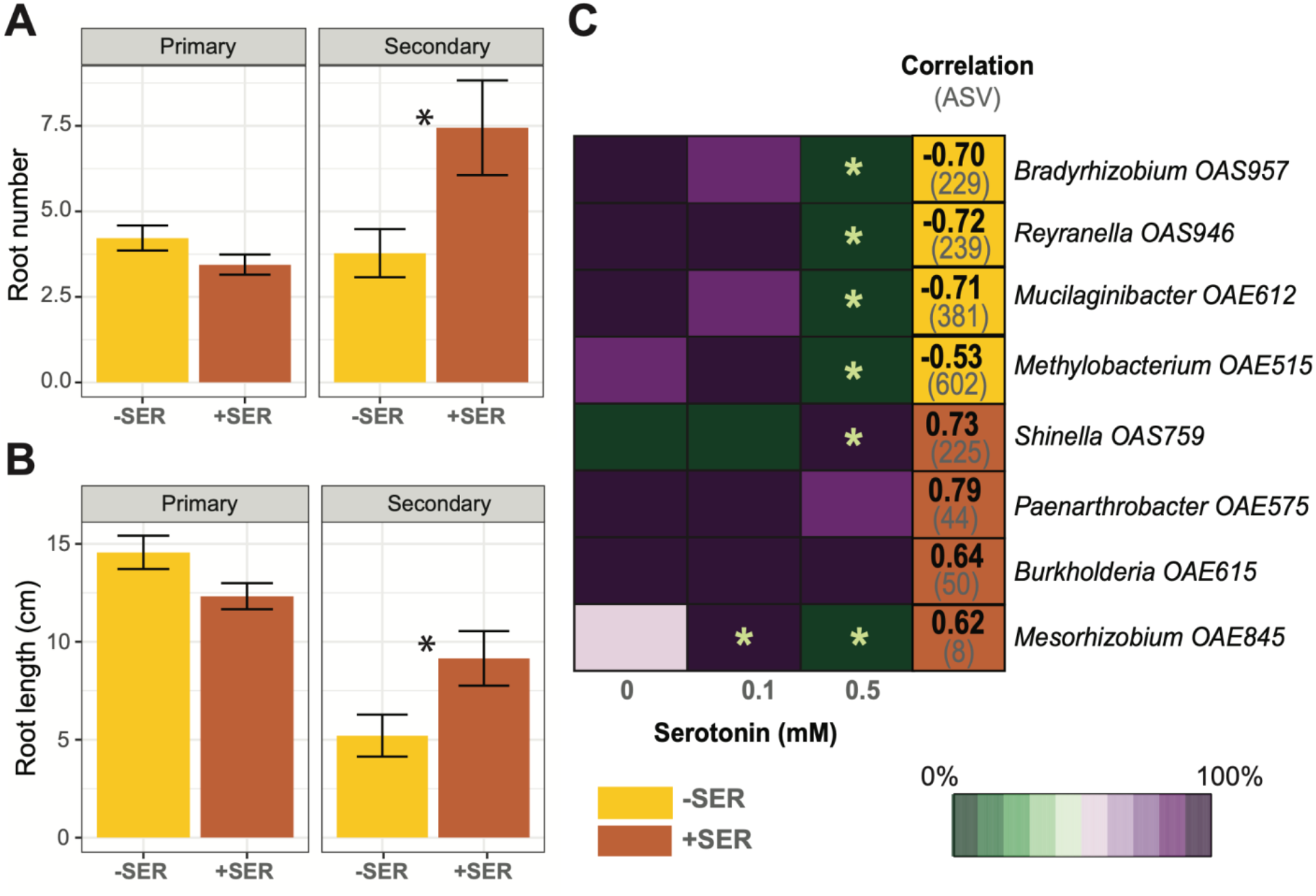
Serotonin effects on switchgrass plant phenotype and growth of rhizosphere microorganisms. (*A,B*) 25 day-old switchgrass seedlings (n=9) grown with exogenous application of 0.1 mM of serotonin (+SER) or controls (−SER). Serotonin effects on secondary roots number (*A*) and total root length (*B*). Significant differences between added-serotonin and controls was assessed by ANOVA, asterisks reflect *P* < 0.05. (*C*) Optical density (OD_600_) of rhizosphere bacteria cultures after 130 hours of growth in 1/10 R2A medium with 0, 0.1, or 0.5 mM of serotonin. Values have been scaled to the highest OD for each isolate across the row. The highest OD of the isolate is 100% (dark purple) and the lowest OD is 0% (dark green), meaning that isolate growth has been completely inhibited. Orange cells indicate isolates related to ASVs with significant negative correlations with serotonin (−SER) and brown cells indicate isolates matched to ASVs with positive correlations (+SER). Positive and negative correlations between specific ASV (shown in brackets) and serotonin shown inside of each cell. Asterisks indicate significantly different OD_600_ between the 0.1 and 0.5 mM serotonin treatments (n=4) and a control treatment without serotonin (0 mM, n=4) at *P* < 0.05 by means of Kruskal-Wallis test.

We also tested serotonin’s effects on eight microbial isolates purified from the switchgrass rhizosphere in marginal soils from Oklahoma (**Fig. 6*C***), and are closely related to the ASVs identified in the switchgrass rhizosphere in this study (≥97% of 16S gene sequence homology). Four of the isolates are closely related to ASVs where we found negative serotonin correlations (’-SER’) and four are related to ASVs with positive serotonin correlations (‘+SER’) (**Fig. 6*C***). Increased serotonin concentrations (0.5 mM) in the isolate growth medium suppressed growth of the -SER isolates based on the OD_600_ readings of this treatment compared to the control (**Fig. 6*C*, Fig. S6**). In contrast, growth of +SER isolates was less sensitive to serotonin additions. Their response varied from slight delays in the lag phase at 0.5 mM of serotonin (*Burkholderia* and *Paenarthrobacter*) to growth stimulation (*Shinella* and *Mesorhizobium*).

## Discussion

### Exometabolites reflect rhizosphere abiotic stress conditions

Plant exudates and microbial metabolites present in the rhizosphere play a significant role in shaping plant-microbe and microbe-microbe relationships under different environmental conditions, including various abiotic stresses (79-82). These rhizosphere metabolites likely consist of a large fraction of the plant exudates, microbial products, and background C compounds present in the bulk soil. A number of studies have analyzed metabolites present in rhizosphere soil of maize, *Arabidopsis*, wheat, rice, wild oat (13, 83-86). However few studies have identified changes of rhizosphere metabolites in soil in response to abiotic stressors. For example, Caddell et al. 2021 investigated the dynamics of sorghum rhizosphere metabolites during drought (87), and Mavrodi et al. (2012) established that phenazines accumulate in the rhizosphere of dryland cereals (88), but responses to nutrient limitation, in particular, have been understudied in rhizosphere soil. Here, we demonstrate that metabolites recovered from the rhizosphere soil of switchgrass shifted in response to nutrient (N, P) and water-limited conditions. We observed three major patterns in switchgrass rhizosphere metabolite shifts: (i) enhanced abundances of aromatic acids when switchgrass was grown in N-limited soil (**Fig. 3*A***); (ii) enhanced abundances of N-containing compounds when N was added (**Fig. 3*C***); and (iii) enhanced abundances of osmolytes in water-limited conditions (**Fig. 3*E***).

Aromatic acids we identified in the rhizosphere of N-limited switchgrass have previously been observed in both plant exudates and rhizosphere soils (12, 89). It has been proposed that the release of this class of metabolites to the environment plays an important role in plant allelopathy and soil phytotoxicity (90), structuring rhizosphere microbiomes (91), suppressing bacterial and fungal pathogens (28, 92) and solubilizing soil P (93). Aromatic acids also contribute to the alleviation of salt stress damage in plants (94) and are involved in quorum sensing and quorum quenching (95). The literature on changes in the abundance and chemical composition of plant root exuded metabolites, particularly aromatic acids, indicate that exudation of these compounds is sensitive to plant developmental stages (12, 13). Recently we found that wild oat exudes the largest quantities of aromatic acids during active growth and that this increase was associated with changes in its rhizosphere microbiome, potentially by attracting beneficial microorganisms (12). These results indicate that plant exudation shifts as the plant’s demands change, for example in response to abiotic stresses. Indeed, multiple studies showed that plants exude compounds that aid in nutrient acquisition when nutrients such as Fe (79, 80), P (79, 81), and N (82) are limited. We observed greater abundances of aromatic acids in our N-limited control soils, consistent with studies that have shown that plants produce more phenolic compounds such as aromatic acids when they are more stressed (93, 95-97) (**Fig. 3*A***). When we alleviated N stress in our system, we observed that metabolites in the switchgrass rhizosphere consisted of fewer aromatic acids and more N-rich compounds, instead (**Fig. 3*C***). Greater exudation of amino acids, nucleosides and other N containing molecules by plants in response to N-amendment has been shown previously (79), and suggests that a higher availability of N enables switchgrass to exude more amino acids, nucleosides, azoles, diazines, and quaternary amines. Furthermore, the increase of N-containing metabolites in the rhizosphere appears to be an indicator of the alleviation of N-stress experienced by switchgrass, given the enhanced root biomass (**Fig. 1*B***) and C availability (43) that resulted from N-addition. Our results indicate that rhizosphere metabolites respond to changes in abiotic stress in a manner consistent with plants and microbes attempting to alleviate abiotic stresses in the rhizosphere.

Consistent with this expectation, we also observed greater production of ectoine, choline, betaine, raffinose and a number of amino acids in water-limited conditions (**Fig. 3*E***). These are widely-known osmolytes - compounds produced by plants and microorganisms to alleviate osmotic stress (98-102). Previous studies have demonstrated these metabolites are more abundant in water-limited soils (99, 103). Notably, some of the same osmolytes were produced in the rhizosphere when water was limiting as when we added N (**Fig. 3*C***). The crossover between the responses to reduced watering and enhanced N availability makes sense given that increased root biomass in the +N and +NP treatments resulted in drier soils relative to controls (**Fig. 1*B,C***) (43). As such, the response of metabolite osmolytes in our system are consistent with osmolytes increasing in abundance when plant and/or microbial cells are experiencing moisture stress in the soil matrix.

### Abiotic stress structures rhizosphere microbiomes

The composition of metabolites in the rhizosphere is defined by a complex mix of plant rhizodeposits, simple molecules from decomposed plant litter, products of microbial catabolism of root exudates and plant polymers, signaling molecules, antibiotics, plant hormones and other active molecules (3, 5, 104-106). Plants are primary contributors to soil C stock and the richest sources of small organic molecules in the soil, releasing 11-40% of photosynthesis-derived carbon into the rhizosphere (4, 31, 104). The metabolites we identified in our study, including sugars, amino acids, organic acids, and nucleosides represent common plant metabolites and have been reported before in exudates of different plants, including grasses and food crops (107). Plants produce and secrete metabolites to modulate the soil environment, and these plant-exuded metabolites are key factors shaping the structure of microbial communities in soil.

The overall community composition that we observed in the switchgrass rhizosphere was consistent with the literature on the taxa found in the rhizosphere of many other grasses and switchgrass-associated soil bacteria (2, 50, 52, 108). We observed that *Actinobacteria* was the dominant switchgrass rhizosphere phylum and *Proteobacteria* (particularly class *Alphaproteobacteria*), *Acidobacteria* and *Verrucomicrobia* were the next-most dominant phyla (**Fig S2*A***). This is in-line with previous research showing that lowland switchgrass genotypes such as the *Alamo* we employed in this study often have increased relative abundances of *Actinobacteria* and *Acidobacteria* (52, 108, 109). Although *Verrucomicrobia* are not generally considered to be “rhizosphere” taxa, Hestrin et al. (2020) reviewed switchgrass microbiome literature and observed that *Acidobacteria, Actinobacteria, Proteobacteria*, and *Verrucomicrobia* were consistently associated with switchgrass roots and rhizosphere soil (110).

Stress-induced changes in metabolite profiles, dominated by the common plant-derived metabolites, lead to the enrichment of specific microbial taxa in the rhizosphere of plants, presumably to assist the plant in counteracting stressors (111). We suggest that switchgrass exudates drive shifts in the rhizosphere metabolome in response to nutrient and water availability, and these changes mediate the assembly of the rhizosphere microbiome. The shifts in rhizosphere bacterial community structure that we observed in response to changes in abiotic stress and in tandem with associated changes in metabolite profiles demonstrate close linkages between these factors and generally support this hypothesis.

The ɑ-diversity of the community was reduced when N was added to the system, which is consistent with numerous prior studies (112, 113). It is hypothesized that alleviating N-limitation can result in the proliferation of bacteria that are adapted for environments with plentiful soil resources at the expense of more diverse taxa better adapted for nutrient-limited environments (114). In terms of taxonomic shifts, we observed the same trend consistently seen by Ramirez et al. (2012) whereby bacterial communities under N-addition have greater abundances of *Actinobacteria* (115). In the same study authors reported a decline in microbial activity (i.e., lower activities of extracellular enzymes) in N-amended soils, suggesting a shift to the preferential utilization of more labile C pools.

In contrast to the lineages that positively responded to N addition, we observed that members of *Verrucomicrobia* and *Acidobacteria* decreased in abundance under N-addition. These slow-growing, oligotrophic lineages have been linked to nutrient deficiencies, in general, and N-limitation, in particular (115), and decreased in abundance in response to more readily available nutrients. These findings corroborate previous field studies, where comparable community changes were observed in the same bacterial groups (115-117). A number of ASVs from *Proteobacteria* also declined in abundance in response to N-addition, including from families such as *Xanthobacteraceae* (ASVs 295, 412, 1189), *Beijerinckiaceae* (ASV 602), and *Burkholderiaceae* (ASV 231, 436, 467, 685) that have been shown to be capable of freely fixing N and making it more available in soil (118, 119). The decline of potential N-fixers in response to N-addition follows from the resource-based economics that govern microbial life history strategies (120). It has also been previously shown that cultivation of perennial grasses increases the abundances of *NifH* gene in the rhizosphere (121), while N-fertilization reduces *NifH* gene abundance in soil from switchgrass stands - though the taxa responsible for the observed decline were likely diverse and not limited to just *Proteobacteria* (122).

We did not observe a strong microbial community response to P addition or watering reduction in our study. The ASVs that were enhanced in prevalence in response to our +P treatment came from a diverse array of lineages with no readily discernible pattern, while those that were decreased in prevalence came primarily from *Actinobacteria*. However, the low number of responsive ASVs (17) makes it difficult to draw inferences. Notably, rhizosphere metabolites in +P treatment were also the lowest compared to other treatments. This could be a result of P dynamics taking longer to affect soil ecosystem dynamics than the single growing season captured in our study, or it could be an indication that P availability is not near as limiting in our system as C or N limitations are. The fewest number of ASVs (6) responded to our -W treatment, with the majority of these also from *Actinobacteria*, though it is notable that no ASVs decreased in prevalence in response to the -W treatment. This may indicate that the dry conditions in these mesocosms made the soil environment more conducive to the growth of certain *Actinobacteria*, these “monoderm” lineages have previously been shown to be drought-tolerant (123).

### Metabolite chemistry associated with suite changes in abundance of specific ASVs

Taxon-specific responses to individual rhizosphere metabolites could be an important driver of rhizosphere bacterial community assembly. Although we emphasize that rhizosphere metabolites are not direct measures of plant exudation, we hypothesized that non-random covariations in the abundances of microorganisms and rhizosphere metabolites across the broad range of abiotic stresses in our treatments could indicate potential functional links between certain metabolites and certain microbial lineages. Our results support this hypothesis, with rhizosphere metabolites shifting with microbial community composition in a similar manner to that observed in the literature for soil microbial communities exposed to changing exudate chemistry.

It has been previously established that plants can exude organic acids in nutrient-limited conditions, at least in part to directly liberate carbon and nutrients into the soil matrix (109). However, it has also been shown that organic acids, and in particular aromatic acids, are exuded by plants as they develop, and that greater abundances of such acids correspond to large-scale shifts in soil microbial community composition (18, 19, 91). Several potential mechanisms of how aromatic acids may modulate rhizomicrobiomes have been proposed, including shifts in soil pH, antimicrobial effects or preferential utilization of these metabolites as a nutrient source by specific microbial taxa geared to decompose their recalcitrant structure (12, 18, 91).

In our association network, we found that half of the six module hubs were organic acids, with chlorogenic acid, an aromatic compound, possessing the most links to microbial ASVs of the three (**Fig. 5**). In addition, we observed that aromatics such as chlorogenic acid, caffeic acid and 4-pyridoxic acid (among others) were most abundant in our control, N-limited marginal soils. These soils also possessed the most diverse microbial communities (**Fig. 1*E***). Reduced concentration of these aromatic acids in our N-amended treatments also corresponded to significantly less diverse rhizosphere bacterial communities, with consistent reductions in similar ASVs (**Figs. 2*B,C***). The ability to metabolize organic acids, in particular, has been linked to the proliferation of taxa in the rhizosphere of a variety of plant hosts (12, 18, 124), which is notable given that organic acids are among the dominant classes of exudate compounds and many plants are known to exude them (along with other rhizodeposits) from their roots during active growth and development (12, 15, 106, 125). Thus, aromatic acids such as chlorogenic acid are likely strong drivers of switchgrass rhizomicrobiome structure.

Two of the three remaining rhizosphere module hub metabolites, serotonin and acetylcholine, have not been extensively studied in the context of soil. Both are produced by plants and microorganisms and are involved in amino acid metabolism. In soil, serotonin can result from the degradation of tryptophan (126), itself a precursor for many essential plant metabolites including plant hormone auxin (53). Changes in the abundances of both acetylcholine and serotonin were also shown to be primarily associated with changes in the N-status of the soil environment (**Fig. 4**).

Interestingly, serotonin was the largest module hub that we observed in our network. In plants serotonin plays important roles in growth, development and response to environmental stresses (53). However, the mechanism of action of this signaling metabolite and its role in the rhizosphere, particularly in plant-microbe interactions, are not well understood. We demonstrated for the first time that serotonin is not only correlated with a large number of ASVs but also impacts plant phenotype and growth of rhizosphere microorganisms. We found that growing switchgrass in microcosms with applications of serotonin increased plant aboveground biomass and significantly enhanced root biomass, the number of secondary roots, and total root length (**Fig. S5 and Fig. 6*A,B***). We found that serotonin can either suppress or stimulate rhizosphere microorganisms and therefore it allows the plant to shape its community to respond to environmental conditions. The established ability to alter root phenotypes combined with the observation of major shifts in abundance of many rhizosphere ASVs in conjunction with the abundance of serotonin and strong impact on microbial growth hint at the possibility that serotonin is a keystone metabolite mediating plant-microbe interactions in the rhizosphere. Recent studies demonstrated that gut microorganisms co-evolved to induce serotonin production by the host and can sense this host-derived serotonin to increase their colonization and fitness in the intestine (54). However, it is also known that many phenylamides, such as serotonin have antibiotic properties (127) which is consistent with our observed suppression of microbial growth and negative correlations between selected microbes and serotonin. In contrast, microbes that have been stimulated by serotonin and corresponded to the ASVs positively correlating with this molecule, possibly could co-metabolize serotonin as a nutrient source or developed a mechanism to detoxify this molecule.

The final module hub metabolite we observed is ectoine, one of the most abundant osmolytes in nature and commonly produced by aerobic heterotrophic bacteria (128). As such, it may be indicative of moisture stress experienced by microbial communities in the rhizosphere. Notably, ectoine abundance was positively associated with *Actinobacteria*, a lineage often thought to be drought-tolerant (129), and associated with ectoine in arid environments (130). While we caution that the mechanisms behind links in a correlation network are difficult to assess, in the context of the broad range of abiotic conditions experienced by the plant and soil in our study, strong correlations between metabolites and microbial ASVs could well imply that they respond similarly (positive links) or differentially (negative links) to these conditions. We note that the organic acids that were identified as module hubs in our association network had mostly positive associations with a diverse array of ASVs, which is supported by the treatment responses we observed for rhizosphere metabolites and microbiome community composition. In contrast, serotonin had negative associations with a diverse array of ASVs, but its few positive associations were almost entirely with *Actinobacteria* lineages, which is supported by the taxonomic responses to N-rich rhizosphere metabolites that we observed. Thus, rhizosphere microbial assembly mediated by metabolites could be important drivers of these covariations - especially when the potential relevance of the chemistry of these compounds to plant-microbial metabolism follows expectations from literature and can be demonstrated in lab settings, as we could for serotonin. State-of-the-art techniques such as stable isotope probing (131), a reductionist approach and experiments in highly controlled environments (132, 133) and improved methods for collection and identification of exudates and plant metabolites from soil systems (131) will provide aid in disentangling links between metabolite chemistry, dynamics or microbial community and plant response to environmental stressors.

## Conclusion

Metabolic changes belowground play a vital role in plant stress resilience and microbial adaptations to environmental change. However, the relationships between rhizosphere metabolite chemistry and the dynamics of microorganisms in soil have been largely overlooked. Our results show that rhizosphere metabolites are sensitive indicators of abiotic conditions in the soil environment that can be linked to the shifts of specific bacterial lineages in response to such changes. Here we show that aromatic acids were enriched in the rhizosphere of N-limited switchgrass and identified microbial lineages associated with this N-limiting condition that were enriched in the presence of these organic acids. In contrast, N-rich metabolites were plentiful in the rhizosphere of N-replete switchgrass, as were fast-growing microbial lineages capable of responding to increased nutrient availability. We contend that the metabolites identified as module hubs in our association network - chlorogenic acid, cinnamic acid, glucuronic acid, serotonin, ectoine, and acetylcholine - merit further study as ‘keystone metabolites’ by structuring soil microbial communities in response to abiotic stress. In conclusion, the rhizosphere metabolite response to nutrient and moisture availability and associated changes in microbiota suggest a putative mechanism of metabolite-driven microbial community assembly under abiotic stress and highlight potential keystone metabolites in the rhizosphere of switchgrass.

## ACKNOWLEDGMENTS

Funding for this project was provided by the United States Department of Energy Office of Biological and Environmental Research (DOE OBER), Genomic Science Program Sustainable Bioenergy award DE-SC0014079 to the University of California–Berkeley, with subawards to Lawrence Berkeley National Laboratory (LBNL), the University of Oklahoma, the Noble Research Institute, and Lawrence Livermore National Laboratory (LLNL, SCW1555). For the metabolite analysis, K.Z., T.N., B.P.B., J.J. and J.S. were supported by m-CAFEs Microbial Community Analysis and Functional Evaluation in Soils, an SFA led by LBNL supported by the U.S. DOE OBER under contract number DE-AC02-05CH11231. We thank David Orme for allowing us to collect soil from his ranch in Anadarko Oklahoma, and Hugh Aljoe, Kelly Craven and Shawn Norton of the Noble Research Institute for facilitating site access and soil collection. We thank Christina Wistrom and the Oxford Tract Greenhouse at the University of California-Berkeley for the infrastructure required to carry out the greenhouse experiment. We thank Katerina Estera-Molina, Anne Kakouridis, Sarah Baker, Steve Blazewicz, Evan Starr, Ka Ki Law, Mengting Yuan, Heejung Cho, Alexa Nicholas, Eoin Brodie, Peter Nico, Caleb Herman, Ashley Campbell, and Amrita Bhattacharyya for their help with the destructive harvest of mesocosms. Thanks also to Madeline Moore, David Sanchez, Cynthia-Jeanette Mancilla and Ilexis Jacoby for their help with processing of soil and nucleic acid samples.

## Notes

### Competing Interest Statement

The authors have declared no competing interest.

