## Supplemental Materials and Methods for "Nutrient and moisture limitation reveal keystone metabolites that link switchgrass rhizosphere metabolome and microbiome dynamics"

### Supplemental Methods

#### DNA extractions

Rhizosphere soil in Lifeguard solution was pelleted for DNA extractions by vortexing samples for 2 min to fully resuspend soil particles before centrifuging for 5 min at 5,000 g and 4 °C, and then re-centrifuging for 2 min after picking out remaining roots, if necessary. The resulting supernatant was discarded and tubes with soil pellets were stored at -80 °C.

Soil pellets were collected by cracking surrounding tubes with a foil-wrapped, DNAse-free hammer on a foil-wrapped, DNAse-free ring stand underlain by dry ice. Pellets were collected in Whirl Paks (Nasco) and stored on dry ice. 0.5 g of each soil sample was aliquoted into a sterile, pre-weighed 2 mL microcentrifuge tube (screw-top, self-standing) after fragmenting the soil pellets in Whirl Paks with the hammer.

Soil DNA was extracted using a modified RNA/DNA phenol chloroform co-extraction protocol via bead-beating (1, 2). Notably, the 5% hexadecyl-trimethyammonium bromide/0.7 M NaCl/240 mM K-PO4 buffer (pH 8) was modified to include 1% β-mercaptoethanol, PEG 8000 was used in place of PEG 6000, and GlycoBlue was used to stain nucleic acid pellets. The combined RNA and DNA in resulting pellets were purified using a modified lithium chloride extraction protocol (3). Purified DNA pellets were stored at -20 °C. DNA was quantified within a week of extraction via PicoGreen fluorescence (4).

Extracted DNA was then fractionated via isopynic centrifugation in a cesium chloride density gradient, using the same method detailed by Blazewicz et al. (5). Immediately following centrifugation, the sample was fractionated into ~40 fractions using a syringe pump. DNA concentrations in each fraction were quantified via PicoGreen fluorescence, and fractions containing DNA were recombined with neighboring fractions to create seven “bins” of equal DNA content (not concentration). Microbial community composition of the DNA in each bin was characterized with a sequencing library prepared at the University of Oklahoma via a phasing amplification technique targeting the V4 region of the 16s ribosomal RNA gene with the 515F and 806R primer set (6) (7). Samples were sequenced on the Illumina MiSeq platform with 2x250 bp format. We note that the process of fractionating the samples before sequencing likely altered the community that we observed - however, this DNA should also be cleaner as a result having been run through the cesium chloride gradient and the results of the sequencing run should contain fewer artifacts.

#### Analysis of 16S rRNA gene sequences pipeline

A total of 10,516,421 raw reads were imported (using the Earth Microbiome Project protocol) into Qiime2, and were demultiplexed, trimmed of non-biological primer sequences, and denoised using DADA2 (225 bp forward reads, 223 bdp reverse) (8). This resulted in 7856 denoised ASVs, which accounted for 3,942,210 (37%) of the total reads. These ASVs were aligned to a rooted tree and assigned taxonomy by a feature classifier. The feature classifier was trained on reads extracted using the aforementioned primer sequences from the SILVA_132 16S reference database for 99% identity. The resulting ASVs were grouped by initial soil sample, such that ASVs from the seven bins the extracted DNA had been fractionated into were re-combined to assess the nature of the switchgrass rhizosphere 16S community and its response to our treatments. These reads and the rooted tree were exported from Qiime2 for further analysis via *phyloseq* in R (9).

In *phyloseq*, all chloroplast or mitochondrial sequences were removed from the dataset, as were bacterial and archaeal sequences that lacked a designation at the phylum level. 7481 ASVs accounting for 3,792,761 reads passed this filter, and were used to calculate α-diversity metrics via the *estimate_richness* function. We tested for significant α-diversity treatment effects using the *lm* and *anova* functions in R, and used the *emmeans* function to test the significance of pairwise comparisons between treatments. For all analyses other than α-diversity, all sequences that only occurred once or twice across the entire dataset were removed in phyloseq. This resulted in 7093 ASVs accounting for 3,792,087 reads, with a range of 104,776 - 194,526 reads per sample. The *distance* function was used to develop a Unifrac dissimilarity matrix by sample that was then assessed for significant treatment effects using the *adonis* function of the *vegan* package in R, with 99999 permutations. We assessed significance of pairwise comparisons between treatments with the *pairwise.adonis* function of the *pairwiseAdonis* package, with 99999 permutations and used the Hochberg correction for multiple comparisons.

Differentially abundant taxa between each of the individual treatments and the control samples were assessed with the *DESeq* function of the *DESeq2* package, employing a local fit for dispersion estimates and optimizing for a significance cutoff of $P < 0.01$ after visually assessing the distribution of P-values. Effect sizes for the magnitude of differences in ASV abundance between the relevant treatment and control samples were calculated as Cohen's *d* in R, using pooled standard deviations between the control samples and those from the relevant treatment.

Cluster analysis of ASVs and metabolite abundances were performed using the *vegan* package in R (10). The most significant associations were selected by filtering (i) differentially abundant ASVs with more than three significant positive or negative correlations (Spearman's rank correlation, $r > 0.7$ or $r < -0.7$, $P<0.05$) with metabolites; and (ii) metabolites with more than one significant positive or negative correlation (Spearman's rank correlation, $r > 0.7$, or $r < -0.7$, $P<0.05$) with ASVs.

**Soil metabolite extraction, analysis and identification**

To extract soil metabolites, 10 ml of ice cold LC-MS-grade water (pH 7.4) was added to 15 ml polypropylene Falcon tubes filled with roots and rhizosphere soil and held on ice. Samples were vigorously vortexed for 10 sec to detach rhizosphere soil from the root. Roots were removed with sterile tweezers and samples were shaken on an orbital shaker (Orbital-Genie, Scientific Industries, Bohemia, NY) at 200 rpm for 1 h at 4 °C then centrifuged at 3220 g for 15 min at 4 °C. For each sample, the supernatant was filtered through a 0.45 µm syringe filter (Pall Acrodisc Supor membrane) into a 15 mL tube. Two ml of supernatant was aliquoted for total organic carbon (TOC) analysis, using a Shimadzu TOC-L Analyzer. Five additional ml of supernatant transferred into a 15 ml Falcon tube, lyophilized using a Labconoco FreeZone 2.5 lyophilizer and stored at -80 °C.

Dried metabolite extracts were resuspended in 1 ml LC-MS grade methanol (Honeywell Burdick & Jackson, Morristown, NJ, USA), vortexed, sonicated on ice for 30 min and incubated at 4 °C overnight. Samples were centrifuged at 3220 g for 15 min at 4 °C, after which the supernatant was dried in a Savant SpeedVac SPD111V (Thermo Scientific, Waltham, MA) for 3 h. Dried samples were resuspended in 100% ice-cold methanol to achieve a final concentration of 3000 ppm/L TOC; these were then sonicated for 15 min using an ultrasonic bath (VWR). Internal standards (1 µg per ml 2-amino-3-bromo-5-methylbenzoic acid, 5 µg per ml $^{13}C$-$^{15}N$-l-phenylalanine and 2 µg per ml 9-anthracenecarboxylic acid) were spiked into the methanol used to resuspend samples. The resulting extracts were filtered with 0.22 µm microcentrifuge PVDF filters (Merck Millipore), and 150 µl aliquots were transferred to LC-MS vials for metabolite analysis.

All chromatography was performed using an Agilent 1290 LC stack, with MS and tandem mass spectrometry (MS/MS) fragmentation data collected in both positive and negative ion mode using a Thermo QExactive mass spectrometer (Thermo Fisher Scientific) in the Northen Lab at Lawrence Berkeley National Laboratory (Supplementary Table S2). For each 3 µl sample injection, full MS spectra were acquired for *m/z* 70–1,050 at 70,000 FWHM (full-width at half-maximum) resolution. MS/MS fragmentation data were acquired using collision energies of 10-40 eV at 17,500 resolution. Sample injection order was randomized and an injection blank of only methanol was run between samples. Normal-phase chromatography was performed using a HILIC column (Agilent InfinityLab Poroshell 120 HILIC-Z, 150 mm × 2.1 mm, 2.7 µm) warmed to 40 °C with a flow rate of 0.45 ml $min^{-1}$ equilibrated with 100% buffer B (95:5 acetonitrile:water w/ 5 mM ammonium acetate) for 1 min, followed by a linear gradient diluting buffer B down to 89% with buffer A (100% water w/ 5 mM ammonium acetate and 5 µM methylene- di-phosphonic acid) for 10 min, then down to 70% B over 4.75 min, then down to 20% B over 0.5 min, and then isocratically held at 20% B for 2.25 min (Table S4).

**Supplemental Figures**

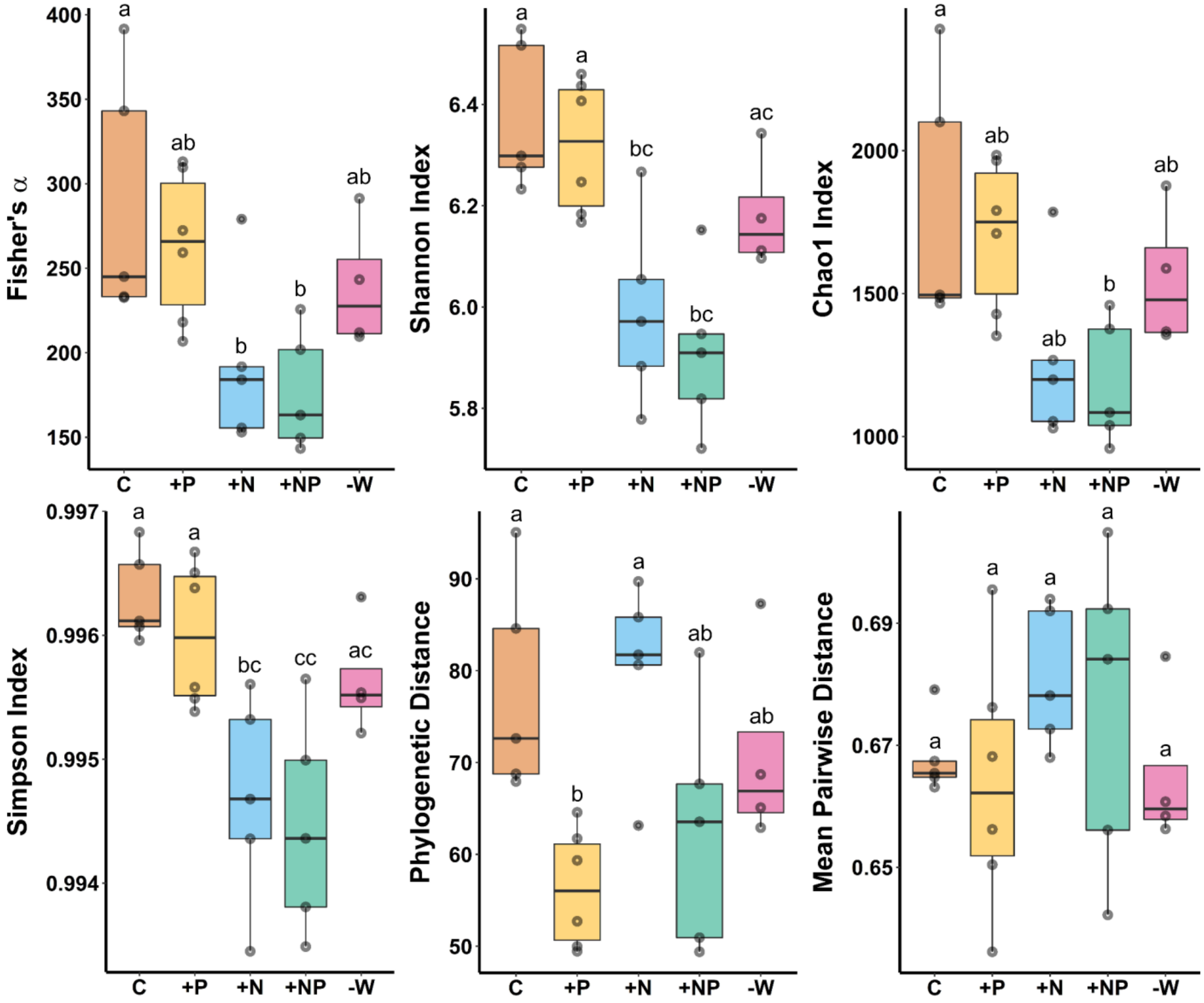


**Fig. S1.** Box-whisker plots (median and 25-75% quartiles) of 16S amplicon sequence variant α-diversity metrics for switchgrass rhizosphere microbial communities grown under five treatments: controls ('C') with nutrient-poor marginal soil, '+P', '+N', and '+NP' mesocosms with phosphorus and/or nitrogen amendments in the top soil horizon, and '-W' mesocosms which received 50% less water relative to the other treatments. Letters represent significantly different post hoc pairwise comparisons via Tukey's test ($P < 0.05$, n=6).

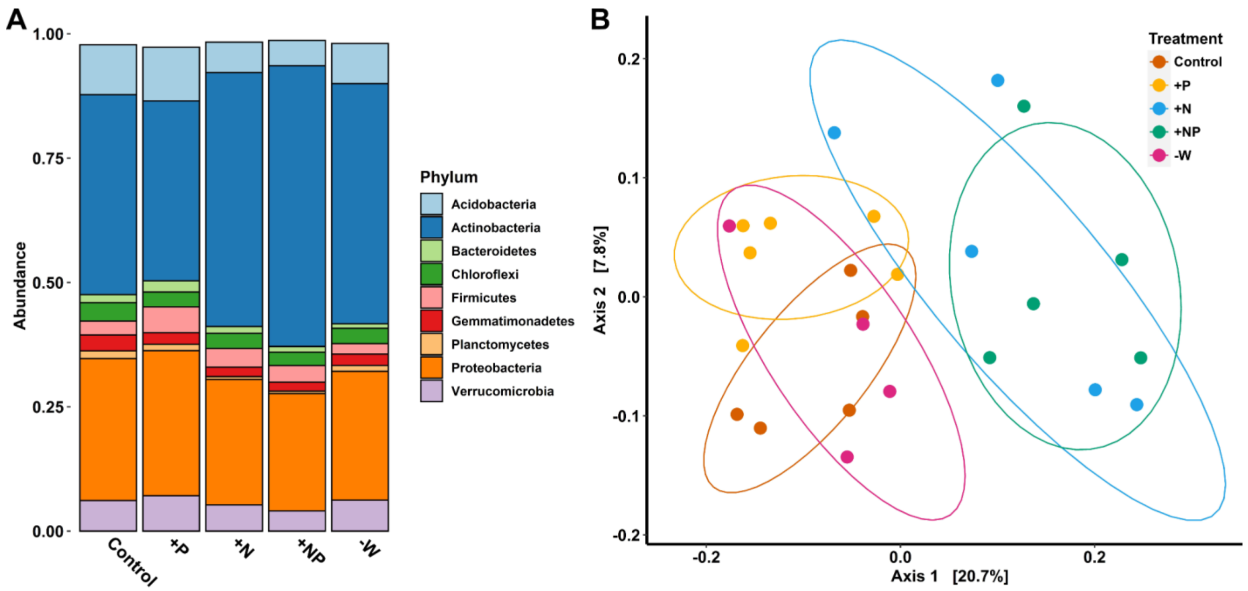


**Fig. S2.** Differences in microbial community composition by treatment, illustrated by (*A*) a barplot of relative abundances of all bacterial phyla with greater that made up more than 1% of the total rhizosphere community and (*B*) principal components analysis of Unifrac distance matrices drawn from the distribution of ASVs in each sample (ellipses indicate 75% confidence intervals).

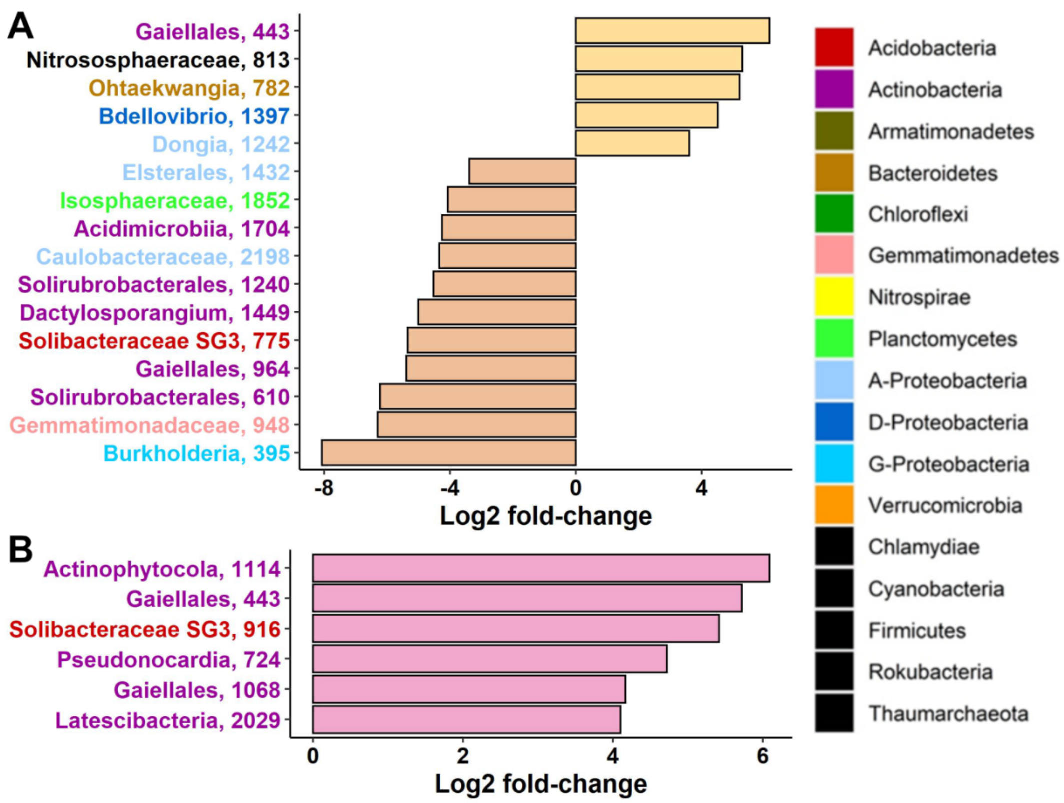


**Fig. S3**. Influence of (*A*) reduced watering and (*B*) phosphorus amendment on switchgrass rhizosphere microbial community structure assessed by DESeq2 analysis (adjusted $P < 0.01$, see **Supplementary Information** for details). ASVs that increased (+ Log2 fold-change) versus decreased in prevalence (- Log2 fold-change) in response to +P or -W treatment are shown. ASVs are presented at the highest available taxonomic resolution, and are colored by class for *Proteobacteria* and by phylum for all other phyla.

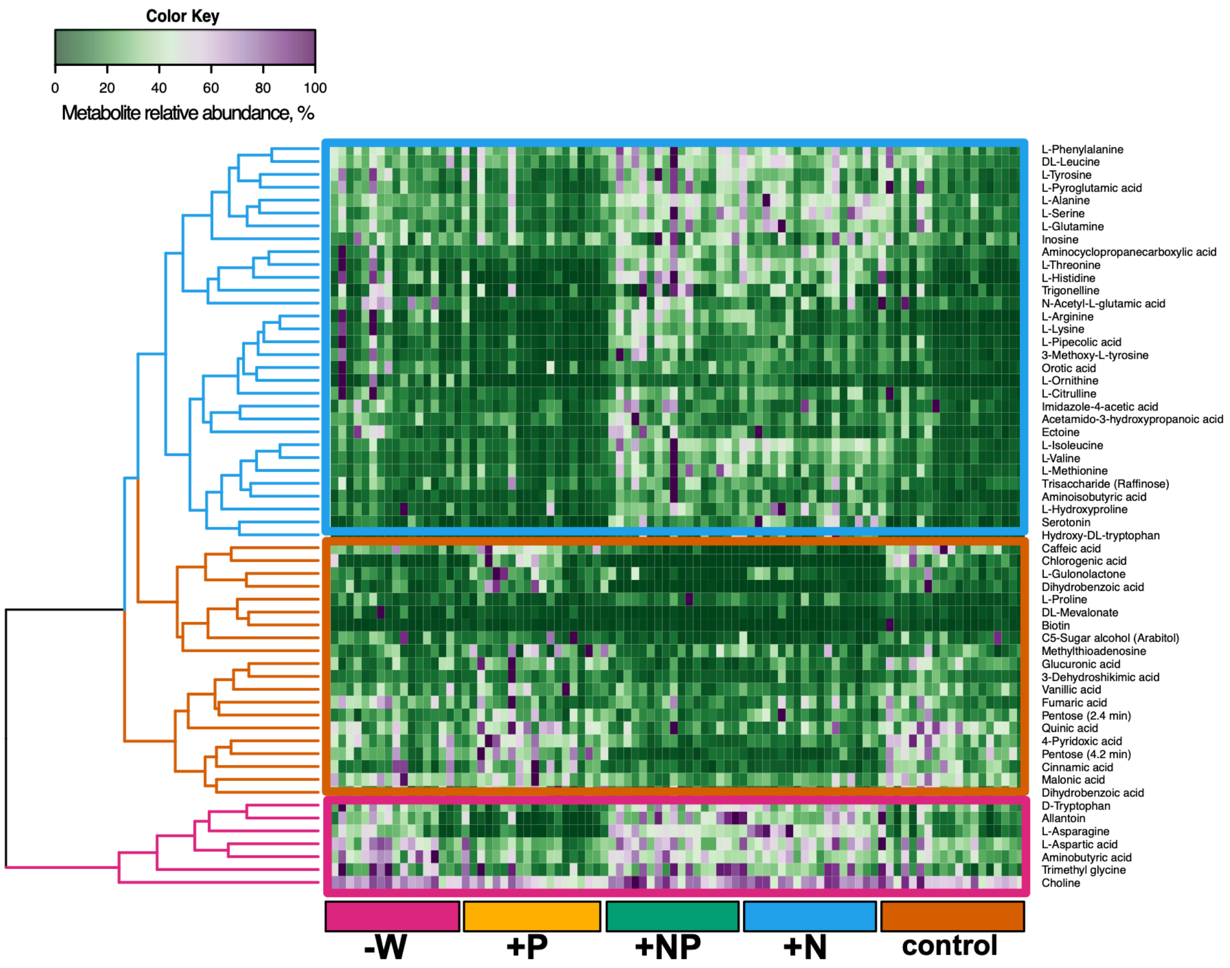


**Fig. S4.** Heatmap of metabolite relative abundances in the rhizosphere of switchgrass grown with five soil nutrient and water treatments. Five treatments include 'control' with nutrient-poor marginal soil, '+P', '+N', and '+NP' mesocosms with phosphorus and/or nitrogen added, and '-W' mesocosms which received 50% reduced water relative to the other treatments. Hierarchical clustering shows three main clusters representing (i) metabolites that were more abundant in +N, +NP treatments (blue lines); (ii) metabolites that were more abundant in no N added treatments (brown lines); (iii) metabolites that were more abundant in water water-limited conditions (pink lines).

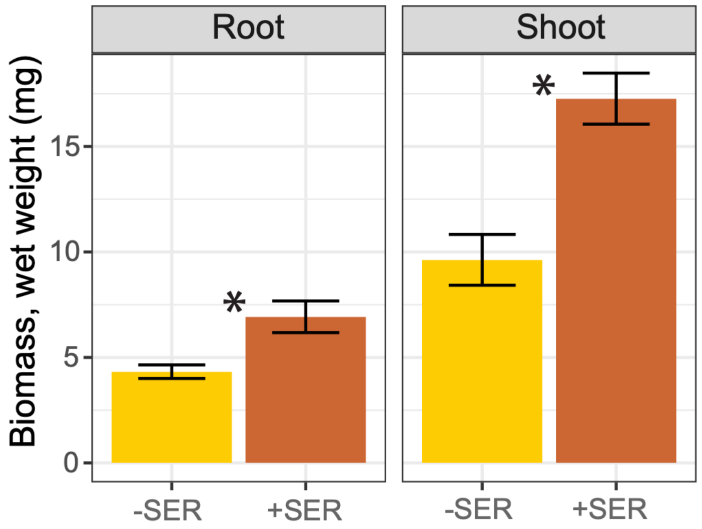


**Fig. S5** Serotonin effects on switchgrass root and shoot biomass of 25 day-old switchgrass seedlings (n=9) grown with exogenous application of 0.1 mM of serotonin or controls. Significant differences between added-serotonin (+SER) and controls (-SER) were assessed by ANOVA; asterisks reflect $P < 0.05$.

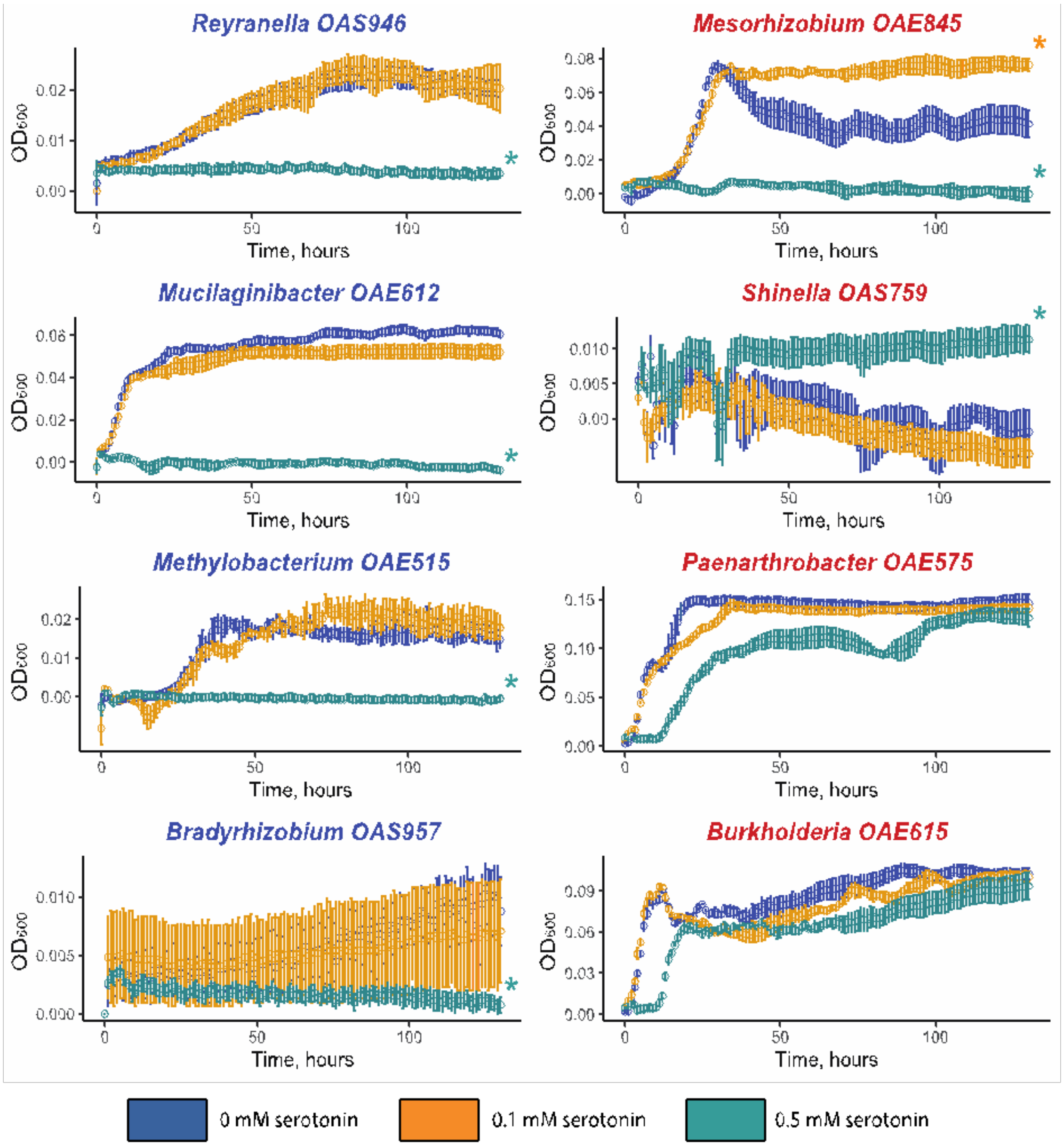


**Fig. S6.** Growth curves of isolates grown in 1/10 R2A medium with 0 (blue), 0.1 (orange), or 0.5 mM (green) serotonin added. 16S rRNA gene sequences of heterotrophic bacteria isolated from marginal fields cultivated with switchgrass, have been compared to ASV sequences formed significant Spearman correlations with serotonin in this experiment (positive and negative). Isolates with a match (E values <1 × $10^{-10}$ and ≥97% of gene sequence homology) to the ASVs with a significant negative Spearman correlation with serotonin are highlighted in blue color and isolates matched to the ASVs with a positive correlation represented in red color. Asterisk indicates significantly different optical density ($OD_{600}$) at 130 hours of isolate growth in serotonin treatments and a control treatment without serotonin (0 mM) at $P < 0.05$ by means of Kruskal-Wallis test. Error bars show the standard error of the mean.
